# Predictive influences on memory encoding: investigating oscillations and the N400 event-related potential component

**DOI:** 10.1101/2024.09.27.615404

**Authors:** Sophie Jano, Matthias Schlesewsky, Alex Chatburn, Scott Coussens, Zachariah R. Cross, Ina Bornkessel-Schlesewsky

**Author notes:** **Corresponding author:** Sophie Jano.

## Abstract

To effectively function in an ever-changing environment, the brain is proposed to make predictions about upcoming information. However, the association between prediction and memory formation and the role of between-subject neural variability in this relationship is unclear. To shed light on the relationship between prediction and memory, the present study reanalysed data from Jano, Chatburn, and colleagues (2024). In the original experiment, participants were exposed to naturalistic images in predictable and unpredictable four-item sequences, after which their memory was tested using an old/new paradigm. In the present analysis (*N* = 46), N400 amplitude and oscillatory power during learning was measured to gauge processes related to prediction error and memory encoding, respectively. This activity was compared with subsequent memory outcomes and individual alpha frequency (IAF) calculated at rest. Linear mixed-effects regressions revealed an alpha power subsequent memory effect that was not related to the amplitude of the N400, suggesting that memory encoding may occur independently of the level of prediction error. Notably, IAF influenced the relationship between theta power, N400 amplitude and subsequent memory, implying that the electrophysiological conditions for successful memory formation differ between individuals. Consequently, the degree to which prediction errors (presumably gauged via the N400) drive memory encoding could depend on inter-individual variability in intrinsic neural activity. These findings emphasise the flexible nature of memory, whilst having potential implications for prediction error-driven accounts of learning.

## Introduction

Perception and cognition may be supported by the brain’s attempts to predict the future (Clark, 2013; Friston, 2005). While predictions could enhance perceptual processing efficiency (Huang & Rao., 2011), they may also incur costs in informational accuracy (as illustrated by their proposed association with false memories; Hubbard et al., 2019; Hubbard & Federmeier, 2024; Jano et al., 2021; Rauss & Born, 2017), although the intricacies of this relationship are currently unclear. According to the predictive coding theory, the degree of departure between what the brain predicts and what it encounters (encoded as prediction error) informs the need for predictive model updating, allowing for more accurate predictions and a consequent reduction of future error (Clark, 2013; den Ouden et al., 2012; Friston, 2005; Rao & Ballard, 1999). As such, prediction errors may prompt memory updating, where the information is encoded into a memory trace (Exton-McGuinness et al., 2015; R. S. Fernández et al., 2016; Mack et al., 2018; Sinclair et al., 2021). On the other hand, predictions are proposed to activate an existing associative memory representation (Bar, 2007, 2009), which may interfere with memory encoding (Sherman & Turk-Browne, 2020). As such, a trade-off between prediction and encoding has been proposed (Hubbard et al., 2019), such that the reliance of predictions on memory retrieval may interfere with the encoding of new information (Sherman et al., 2022; Sherman & Turk-Browne, 2020). This could manifest in a tendency for shallower processing of predictable stimuli, where the brain verifies congruence with predictions rather than meticulously processing each instance of the sensory information (Hubbard et al., 2019; Van Berkum, 2010). Consequently, information that is surprising or incongruent with expectations should be better remembered than predictable information; a suggestion that is supported by prior research (e.g., Federmeier et al., 2007; Greve et al., 2017; Rouhani et al., 2018; Worthen & Roark, 2002). However, other literature demonstrates enhanced memory for predictable information (e.g., Gronau & Shachar, 2015; Turan et al., 2023), and for high frequency stimuli (G. Fernández et al., 1998). This highlights the potential complexity surrounding the relationship between prediction and memory, which may be difficult to capture with behavioural analyses alone. As such, an investigation into the effect of prediction on memory at the neurophysiological level may extend upon existing behavioural research (e.g., Gronau & Shachar, 2015; Ortiz-Tudela et al., 2023), as processes relating to prediction and memory may be gauged more explicitly.

### The relationship between oscillations, memory, and prediction

Oscillatory activity can inform the nature of neuronal communication and information representation within the brain (Buzsáki & Draguhn, 2004; Schnitzler & Gross, 2005). Particularly relevant are theta oscillations in the approximate 4–7Hz range, which have strong ties to memory (e.g., Hasselmo, 2005; Hasselmo & Stern, 2014), prediction error processing (e.g., Cavanagh et al., 2010), and to broader functions such as spatial navigation (Buzsáki, 2005). Increases in theta power during learning have been associated with enhanced subsequent memory at test, highlighting the relevance of this activity for understandings of memory encoding (Hanslmayr et al., 2009; Klimesch, Doppelmayer, et al., 1996, Klimesch et al., 1997; White et al., 2013). Theta power is typically prominent over frontal regions (Hanslmayr et al., 2009; Klimesch et al., 1997; White et al., 2013), prompting the suggestion that scalp-recorded frontal midline theta (FMT) is especially relevant for successful episodic memory encoding (for a review see Hsieh & Ranganath, 2014). Mechanistically, the relationship between theta oscillations and memory functioning could be supported by theta’s ties to long-term potentiation (LTP; a type of synaptic plasticity involving an enduring strengthening of synaptic potentials; Collingridge et al., 2010; Leung & Law, 2020; Martin et al., 2000). LTP is proposed to underly memory formation (Martin et al., 2000), and to be elicited during phases of hippocampal theta that reflect depolarization (for a review, see Leung & Law, 2020). Consequently, theta rhythms may be integral for the synaptic changes that give rise to memory encoding (Leung & Law, 2020), suggesting that such activity can provide insight into the extent to which predictions facilitate or impair memory.

Not only are theta rhythms relevant for memory processing, but they have also been heavily studied in the prediction error literature. Prior research linking theta oscillations to prediction demonstrates greater theta power (i.e., increased synchrony; Klimesch, 1999) in response to incongruent information (Dini et al., 2022), semantic violations (Pu et al., 2020), prediction errors during reinforcement learning (Cavanagh et al., 2010; Mas-Herrero & Marco-Pallarés, 2014), and behavioural errors (Cohen, 2011). As prediction errors result from a discrepancy between predicted and actual events (Friston, 2005; Hohwy, 2020), a behavioural error would likely also induce a prediction error, highlighting the functional relevance of theta power for predictive processing. The link between theta and error processing is also supported by its relationship to the error-related negativity (ERN); an event-related potential (ERP) component that peaks after an erroneous response (Holroyd & Coles, 2002; Luu et al., 2004; Trujillo & Allen, 2007). Moreover, theta power increases following negative feedback have been shown to predict subsequent behavioural adjustments (Van de Vijver et al., 2011), suggesting that theta oscillations play an important role in the memory encoding processes related to prediction error correction and learning. From this perspective, if predictions impair memory encoding, theta power should theoretically be weaker when individuals are predicting, and stronger in the face of a prediction error, indicating enhanced encoding.

While memory formation may be fostered by increased power in the theta frequency range, it may also be supported by changes in power in the alpha oscillatory band (∼8–12 Hz; Hanslmayr et al., 2009, 2012). Desynchronisation in the alpha rhythm during learning has been linked to enhanced memory at test (Hanslmayr et al., 2009; Klimesch, Schimke, et al., 1996; Klimesch et al., 1997; Sederberg et al., 2003, for a review see Hanslmayr et al., 2012), suggesting that alpha power is functionally relevant for memory processing. Klimesch and colleagues (2007) postulate that alpha is linked to inhibition, such that greater alpha synchronisation (giving rise to stronger alpha power; Klimesch, 1999) reflects increased inhibition, while alpha desynchronisation reflects a discharge of this inhibitory mechanism. The relationship between alpha power, prediction and memory was explored and interpreted in the context of inhibition by Ergo and colleagues (2019), who manipulated the degree of reward prediction error (RPE) and found enhanced recognition for large, positive RPEs (where the reward is better than expected). Supplementing these behavioural findings was the observation that alpha power decreased following positive RPEs and increased following negatively valanced RPEs (when the reward is worse than expected). This increase in response to negative RPEs was proposed to reflect the inhibition of task-irrelevant stimulus representations (Ergo et al., 2019). In contrast, positive RPEs could have facilitated memory encoding (evidenced by alpha power decreases) as individuals learnt the contingencies that led to positive outcomes. This interpretation implies that prediction errors can at times impair memory, somewhat challenging traditional predictive coding schemes where prediction errors drive model correction (Rao & Ballard, 1999).

Although the literature described above suggests that memory encoding is supported by decreases in alpha power, Klimesch and colleagues (2007) conversely propose that increases in alpha power prompt the suppression of memory retrieval mechanisms, facilitating memory encoding. Consequently, alpha desynchronisation may be linked to memory retrieval (Klimesch et al., 2007), suggesting that alpha power could be weakest during prediction (when a prior memory representation is likely retrieved; Bar, 2007, 2009; Schacter et al., 2008), and strongest during processing of unpredictable stimuli. While this highlights the complex relationship between alpha power, inhibition, and memory performance, it also emphasises the importance of alpha oscillations for understanding the influence of prediction on memory uptake. This may be further supplemented by an analysis of ERP activity, which can provide a fine-grained, real-time measure of the processes associated with prediction (Luck, 2014).

### The importance of the N400 event-related potential (ERP) for understanding the effect of predictive processing on memory

Unlike oscillations, which fluctuate across longer time periods, ERPs reflect transient activity that can be captured with precise temporal accuracy (Luck, 2014). As such, ERPs are important for measuring nuanced changes in cognition as expectations unfold. Specifically, an analysis of the N400 ERP could allow for learning to be tracked as predictions naturally develop in the absence of explicit behavioural responses or feedback. The N400 peaks approximately 400ms post-stimulus onset and is sensitive to the degree of stimulus expectancy, exhibiting larger (more negative) amplitudes to unexpected items (for a review, see Kutas & Federmeier, 2011). Whilst this pattern has primarily been observed in language paradigms (e.g., Federmeier & Kutas, 1999; Kutas & Hillyard, 1980, 1984), N400 effects are also observable in non-linguistic contexts, linking the component to meaning processing more broadly (Amoruso et al., 2013; Federmeier & Kutas, 2011; Sitnikova et al., 2008; Urgen et al., 2018). The correlation between the N400 and stimulus predictability in turn supports its proposed relationship with prediction error, where the magnitude of the N400 reflects the extent of prediction violation (e.g., Bornkessel-Schlesewsky & Schlesewsky, 2019; Eddine et al., 2024; Hodapp & Rabovsky, 2021; Rabovsky & McRae, 2014). According to the predictive coding theory, the brain reduces redundancy by representing unexplained sensory data in the form of a prediction error (Feldman & Friston, 2010; Huang & Rao, 2011; Rao & Ballard, 1999). Consequently, when sensory information is explained, it may not be represented (Rao & Ballard, 1999). This may render a prediction error-related component (such as the N400) a useful measure of predictive processing in the brain. As such, an analysis of the N400, due to its ties to prediction error (e.g., Bornkessel-Schlesewsky & Schlesewsky, 2019), could be effective in capturing prediction more directly than, for example, pre-stimulus neural activity at the time of prediction (as we observed in a previous study; Jano, Cross, et al., 2024).

Moreover, the question remains as to whether oscillatory patterns and ERPs reflect the same or differing processes in the context of prediction. For example, whilst theta activity has been considered an error processing signal in the past, being proposed to underlie ERPs such as the ERN and the feedback-related negativity (FRN; Cavanagh et al., 2010; Cavanagh & Frank, 2014; Trujillo & Allen, 2007), its general relationship to memory (described above) could also suggest that it reflects the updating that accompanies an error signal, as opposed to the error itself. This is supported by Munneke and colleagues’ (2015) study, which suggests that theta/delta power (in the approximate 2–8Hz range) and the ERN reflect both disparate but complementary processes. Prior research has also observed increased N400 amplitudes to incongruent stimuli alongside oscillatory theta increases (Dini et al., 2022), highlighting the strong connection between these neurophysiological measures. However, few studies directly compare the relationship between oscillations and N400 patterns during learning with behavioural subsequent memory outcomes at test. While a previous study by Klimesch, Schimke, and colleagues (1996) examined the N400 and the oscillatory rhythms that contribute to later remembering, the direct relationship between these combined factors was not explicitly tested. Additionally, most studies, if engaging in a combined oscillatory and ERP investigation, measure the N400 (and related components, such as the FN400) at test to gauge the level of memory recognition (e.g., Chen & Caplan, 2017; Ordin et al., 2020). Consequently, a comparison of N400 patterns with oscillatory activity during learning may provide insight into the extent to which predictability (measured via the N400) influences memory encoding (potentially gauged via oscillatory rhythms). Relating this to memory recognition could in turn shed light on how these neurophysiological components influence later memory outcomes, potentially capturing variability that extends beyond the behavioural domain.

### Individual alpha frequency (IAF)

While a combined investigation of oscillatory patterns, N400 activity and subsequent memory may be useful for understanding the relationship between prediction and memory, supplementing this with an analysis of individual neural variability could provide insight into the individual variability associated with prediction. In a previous study (Jano, Chatburn, et al., 2024), we observed a positive relationship between the magnitude of the N400 during a prediction task and later memory recognition (for similar effects relating to implicit memory see Hodapp & Rabovsky, 2021). However, this trend was only observed for individuals with a low individual alpha frequency (IAF; the dominant frequency within the range of the alpha band; Corcoran et al., 2018; Klimesch, 1999; Jano, Chatburn, et al., 2024). This raises the question of whether prediction errors prompt memory updating for individuals with a high IAF, or whether memory encoding occurs independently to a prediction error signal for these individuals. Furthermore, the relationship between IAF and prediction is currently unclear, with some research suggesting that individuals with a high IAF have a stronger propensity for model updating (Kurthen et al., 2020; particularly under conditions of high prior stimulus surprisal; Jano, Cross, et al., 2024), and other research observing greater model updating for low versus high IAF individuals (Bornkessel-Schlesewsky et al., 2022). Given also that IAF is positively correlated with general intelligence (Grandy et al., 2013), and with memory outcomes (Klimesch et al., 1990, 1993), it is important to explore its relationship to prediction and memory formation in more detail.

### The present study

The current study sought to further examine the effect of prediction on memory via a reanalysis of Jano, Chatburn, and colleagues (2024), with a focus on oscillations, the N400, and individual neural variability (as measured via IAF). In our original experiment, participants took part in a sequence learning task, where they were presented with predictable and unpredictable sequences of naturalistic outdoor scene images, and their memory for the images was subsequently tested (Jano, Chatburn, et al., 2024). Importantly, memory performance was not quantified on an item-by-item level but was instead measured at the aggregate level via the sensitivity measure d’ (McNicol, 2004). However, a subsequent memory analysis measuring hits and misses at test, coupled with an investigation into oscillatory patterns during learning, may provide a more granular and nuanced understanding of how neural patterns during prediction influence the encoding of a particular stimulus into memory.

Based on the relationship between theta oscillatory activity and subsequent memory described in previous literature, it was hypothesised that: a) theta power would be stronger for images in unpredictable versus predictable sequences. As the research linking alpha rhythms to inhibition and subsequent memory is mixed, we also formulated a non-directional hypothesis stating that b) alpha power would differ during the processing of predictable versus unpredictable images. It was further predicted that: c) theta power during learning would be greatest for subsequently remembered items (memory hits) as compared to subsequently forgotten items (memory misses), while d) alpha power during learning would be associated with memory recognition performance at test. Importantly, while the previous study included a categorical manipulation of stimulus predictability, changes in the N400 were proposed to reflect more subtle, item-specific differences in predictability and potentially, the level of prediction error elicited by a stimulus (Jano, Chatburn, et al., 2024). As such, we also aimed to consolidate the distinct perspectives that separately link prediction to the N400 ERP, to oscillations, and to memory outcomes, hypothesising that e) N400 patterns would relate to oscillatory activity during learning, and that this would be associated with later remembering (quantified as hits and misses). Finally, we also aimed to capture the inherently individual nature of prediction via the inclusion of resting state individual alpha frequency (IAF) in the above analyses.

## Methods

### Original study

The present reanalysis used data from our previous study (Jano, Chatburn, et al., 2024; https://osf.io/qvjer/), which included 48 healthy individuals in the final sample (mean age = 26.8 years old, SD = 6.6 years). Participants completed a task adapted from Sherman and Turk-Browne (2020). They first underwent a learning phase where they viewed 102 four-item ABCD sequences of outdoor scene images, belonging to 12 categories of scenes. Sixty-eight of these sequences were predictable, such that images from the same categories in the same order were presented. The remaining 34 sequences contained images from other categories in a random order, such that multiple images from the same category could be present in the sequence, rendering it impossible to predict the upcoming image category. Each item was unique and was only viewed once. Images were presented for 1000ms and were separated by a 500ms inter-stimulus interval displaying a fixation dot in the middle of the screen. Each sequence was followed by a 2000ms interval where participants viewed the same fixation dot as in the interstimulus interval. To ensure that participants were attending to the task, a prompt would randomly appear in the between-sequence intervals, instructing participants to press the ‘Enter’ key. This occurred on seven occasions, and participants were required to press the key as quickly as possible. Participants were told that this was a secondary task, and that their primary objective was to pay attention to the images in the sequences. Individuals were not informed of the relationships between the images and were not asked to actively predict. This is because we sought to probe naturalistic predictive processing, which should be engaged in scenarios where statistical regularities are present (Baker et al., 2014; Bar, 2007; Mumford, 1992). Participants’ electroencephalography (EEG) was recorded during the task using a 32-channel Brainvision ActiCHamp system (Brain Products GmbH, Gilching, Germany).

To avoid recency effects, participants then engaged in a 15-minute digit span task, after which they underwent a surprise memory recognition test. Old images that were seen during the learning phase, along with categorically related lures that had not been viewed during learning were presented in a random order, external to any sequential relationships. Participants were instructed to make an old/new judgement to each image (‘old’ being that they remembered seeing the image during learning, and ‘new’ meaning that they did not remember the image). A keypress was used to indicate their response (z = old, m = new), allowing for the measurement of participants’ memory recognition performance. For a full description of the study materials, protocol, and EEG recording procedure, see Jano, Chatburn, and colleagues (2024), and for a schematic of the task, see *Figure 1*.

**Figure 1.**
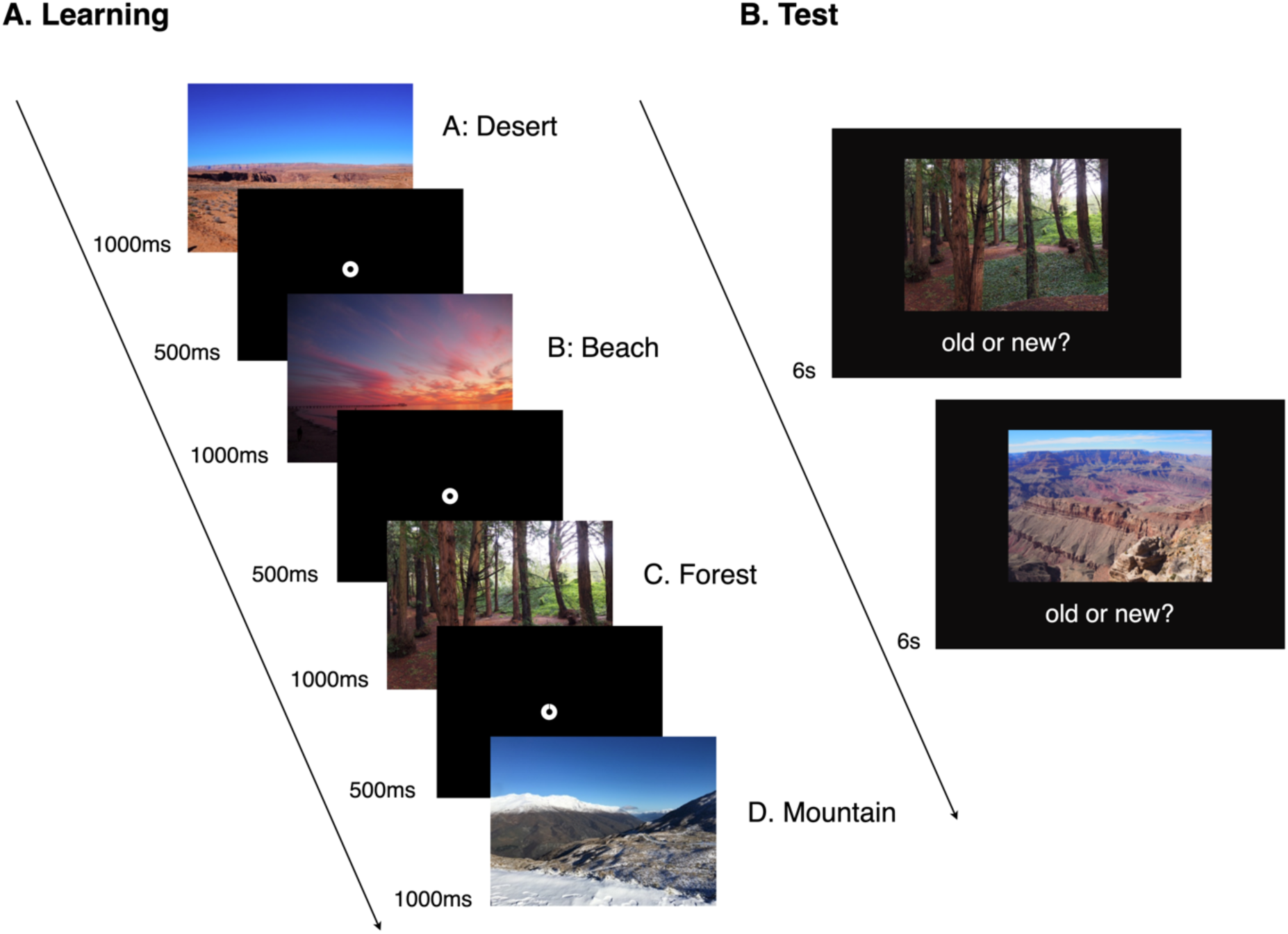
Adapted from Jano, Chatburn, and colleagues (2024) showing the outdoor scene task. **(A)** Learning phase: Participants were instructed to attend to four-item sequences of outdoor scene images in predictable and unpredictable relationships. **(B)** Recognition test: Participants were presented with old images seen during learning, and lure images that were not encountered during the learning phase. Participants were tasked with making an old/new judgement for each image.

### Participants

The sample for the present reanalysis included a total of 46 participants (mean age = 26.5, SD = 6.5, 15 males, 31 females). The sample size was slightly reduced from the previous experiment (which included 48 participants) as IAF metrics for two individuals could not be estimated. As time frequency measures were obtained by adjusting frequency bands according to each individual’s IAF, oscillatory bandwidths could not be defined for these participants, and they were hence removed. Participants’ ages ranged from 18-39, and all individuals were right-handed with corrected-to-normal vision. They also did not have any diagnosed psychiatric conditions, intellectual impairments, or a diagnosis of dyslexia. Finally, they were not taking medication that could affect the EEG and had not taken recreational drugs in the previous six months.

## Data analysis

### Electroencephalography (EEG) data processing

The data from Jano, Chatburn, and colleagues (2024) was used for the current analysis. Raw EEG data were re-referenced to mastoid channels (TP9 and TP10) and Fp1 and Fp2 were set as electrooculogram (EOG) channels. An independent component analysis (ICA) was fit to a 1 to 40Hz band-pass filtered copy of the data, before subsequently being applied to the raw data to detect and remove ocular artefacts. Following this, bad channels were interpolated, and the data were band-pass filtered from 0.1 to 30Hz. Next, 5 second epochs were generated from the pre-processed EEG data around each image in the sequences using MNE-Python version 1.6.1 (Gramfort et al., 2013). Epochs ranged from -1 to 4 seconds surrounding each image to avoid edge artifacts and to capture a sufficient number of oscillatory samples. Finally, the *Autoreject* function (Jas et al., 2017) was run, which interpolated or excluded bad epochs. This resulted in an average of 249.8 epochs remaining in the predictable condition and 124.5 remaining in the unpredictable condition (total images in the predictable condition = 272, total images in the unpredictable condition = 136).

### Metrics from the original study

The IAF data and the N400 ERP data from the original study were used in the present experiment. For a full description of how these metrics were calculated in the original analysis, see Jano, Chatburn, and colleagues (2024). To briefly summarise, IAF was computed across channels P3, P4, Pz, Oz, O1 and O2 from eyes-closed resting state recordings that were captured at the start of the experiment. The *Philistine* package (Alday, 2018) was used to calculate participants’ IAF within 7–13 Hz. Centre of gravity (CoG) was specifically used as a measure of IAF, which reflects the mean power within the alpha frequency band (Corcoran et al., 2018). These procedures consequently resulted in single-subject IAF values that were used for the current reanalysis.

Mean N400 ERP activity across all channels was computed over 1.3 second epochs, across a 300-500ms window following stimulus onset (in line with prior research; Federmeier et al., 2007; Hodapp & Rabovsky, 2021; Hubbard et al., 2019; Van Berkum et al., 2003). Activity in a pre-stimulus window (−300 to 0ms) was also generated, to obtain an ERP baseline estimate for analyses in which N400 activity was included as a predictor. Following ERP generation, the data were averaged over frontal channels (Fz, F3, F7, FT9, FC5, FC1, FT10, FC6, FC2, F4, F8), in line with previous research (Kutas & Federmeier, 2011; Urgen et al., 2018), and to maintain consistency with the original study (Jano, Chatburn, et al., 2024). This resulted in mean ERP amplitude data per subject and item, for both the N400 and the prestimulus window. For grand average plotting, downsampled data (at 125Hz) spanning the full epochs were used and plots were generated using the *ggplot2* (Wickham, 2016) package in R version 4.3.1.

### Time frequency analysis

A time frequency analysis was performed using MNE-Python version 1.6.1 (Gramfort et al., 2013). To measure the oscillatory activity associated with each image during the learning phase, frequency bands were first individually adjusted according to each participant’s IAF (specifically, their CoG values). This procedure is recommended by Klimesch (1999) and Klimesch and colleagues (1997). Once the upper and lower bounds of the alpha and theta frequencies were defined for each participant, a time frequency analysis was performed on the 5 second epochs, on a trial-by-trial basis using the Morlet wavelet transformation from the *tfr_morlet* function. To define the length of the Morlet wavelet, the data within each frequency band was divided into five points, and the number of cycles was calculated as each frequency value divided by four. A fast Fourier transform (FFT) was applied using the *use_fft = True* argument. Following this, the data were initially extracted over five 200ms long windows across the duration of image presentation (0-1000ms), to capture transient alterations in oscillatory activity that may occur within each segment. However, the inclusion of such windows in statistical models led to multicollinearity, suggesting that the windows explained similar variance in the data. As such, we opted to instead collapse the data over one window of interest (200ms to 1000ms post-image onset). This window was chosen to avoid activity relating to early sensory processing (Pratt, 2011). It was also based on similar research, where alpha and theta oscillations associated with error processing and/or subsequent memory appeared maximal within an approximate 200ms to 700ms window (e.g., Hald et al., 2006; Klimesch, Schimke, et al., 1996; Van de Vijver et al., 2011). For the present study, the upper limit was extended to 1000ms to account for the complexity of the naturalistic outdoor scene images, and to capture the whole duration of image presentation. Additionally, a baseline window ranging from -350ms to -150ms prior to image onset was defined, to capture the time-frequency activity in the inter-stimulus window whilst avoiding overlap with the previous stimulus and the upcoming stimulus. Activity within each window was then averaged across time points, resulting in a mean amplitude measure per window and frequency band of interest, for each item and channel. To perform a power normalization, a decibel conversion was then applied to the time frequency data using the below formula:

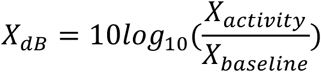

### In-task 1/f aperiodic activity

Oscillations reflect brain activity in the periodic domain, presenting as peaks in amplitude at distinguishable temporal frequencies (He, 2014). Such patterns are purportedly confounded with the aperiodic (scale-free) component, which is independent of specific frequency bands, and which follows the 1/*f* power law (Donoghue, Dominguez, et al., 2020; Donoghue, Haller, et al., 2020; He, 2014; He et al., 2010). Consequently, recent accounts emphasise the importance of measuring both periodic and aperiodic activity during neurophysiological investigations, to estimate effects relating to oscillatory components more directly (Donoghue, Dominguez, et al., 2020). The present analysis employed the Irregular Resampling Auto-Spectral Analysis (*IRASA;* version 0.6.4) function (Wen & Liu, 2016) implemented through the YASA toolbox (Vallat & Walker, 2021) to separate the aperiodic component from the periodic component. The function was applied to the same 5-second epochs used in the time frequency analysis (described above), across a frequency range of 1 to 30Hz, with a sliding window length of 2 seconds. This returned the fractal aperiodic values for each channel and event of interest. Importantly, whilst this method also produced measures of the aperiodic-independent oscillatory activity, this data could not be segmented into windows of interest as it did not return information at the temporal level that could be subsequently sliced. An alternative may have been to compute the function over shorter windows. However, we sought to estimate the spectrum over longer epochs to avoid edge artifacts, and to ensure that a sufficient number of samples was obtained. As such, only the aperiodic 1/*f* slope values from this analysis were used for subsequent investigation. This data was included as a *covariate* in statistical models with oscillatory power calculated from the *TFR_morlet* function (described above). This allowed for the aperiodic patterns related to the events of interest to be accounted for.

### Subsequent memory

Behavioural performance on the memory recognition test was quantified on an item-by-item basis via a subsequent memory analysis, based on participants’ ‘old’ and ‘new’ responses to stimuli. ‘Old’ responses to old images seen during the learning phase were labelled as subsequently remembered ‘hits,’ whilst ‘new’ responses to old images were defined as subsequently forgotten ‘misses.’ As we were primarily interested in how activity during learning predicted encoding and subsequent remembering, the analysis did not focus on behavioural responses to new lure items.

### Statistical analysis of hypothesized effects

Statistical analysis of the data was performed using linear mixed-effects models (LMMs) computed using the *lme4* package (Bates et al., 2014) in R version 4.3.2. These models allowed for estimation of the effects of interest, whilst accounting for random variation. For models predicting theta power, data were summarised across electrodes F3, Fz and F4, as prior research suggests that mid-frontal theta activity is relevant for memory processing (Hsieh & Ranganath, 2014). For analyses predicting alpha power, a more exploratory, data-driven approach was first taken, such that electrodes were grouped according to sagitallity (anterior, central, posterior) and averaged, resulting in a mean value per level. Sagitallity was then entered as a predictor into the models.

All continuous predictors in the LMMs were scaled and mean-centred, whilst categorical predictors (image condition [predictable, unpredictable], image position [A, B, C, D], recognition [hit, miss] and sagitallity [anterior, central, frontal]) were sum contrast coded with the *contr.sum* function from the *car* package (Fox & Weisberg, 2019). All models included random effects on the intercept of subject and item, whilst random slopes of predictors were included when this did not impact model convergence or lead to singular fits (for further discussion of the implications of model overcomplexity, see Bates et al., 2018). To account for task-related aperiodic patterns, on-task 1/*f* slopes were included in all models as a covariate with an additive term. Additionally, whilst image position was not a primary factor of interest, it was included in models 1 and 4 to account for the structure of the underlying data. However, as this structure was accounted for by the variable ‘recognition’ in the other models, position was omitted as a predictor to reduce unnecessary complexity (for a full specification of final statistical model structures, see *Table 1*). P-values were obtained via type II Wald tests from the *car* package (Fox & Weisberg, 2019). Model effects were estimated using *ggeffect* (Lüdecke, 2018), and were visualised using *ggplot2* (Wickham, 2016).

**Table 1.**
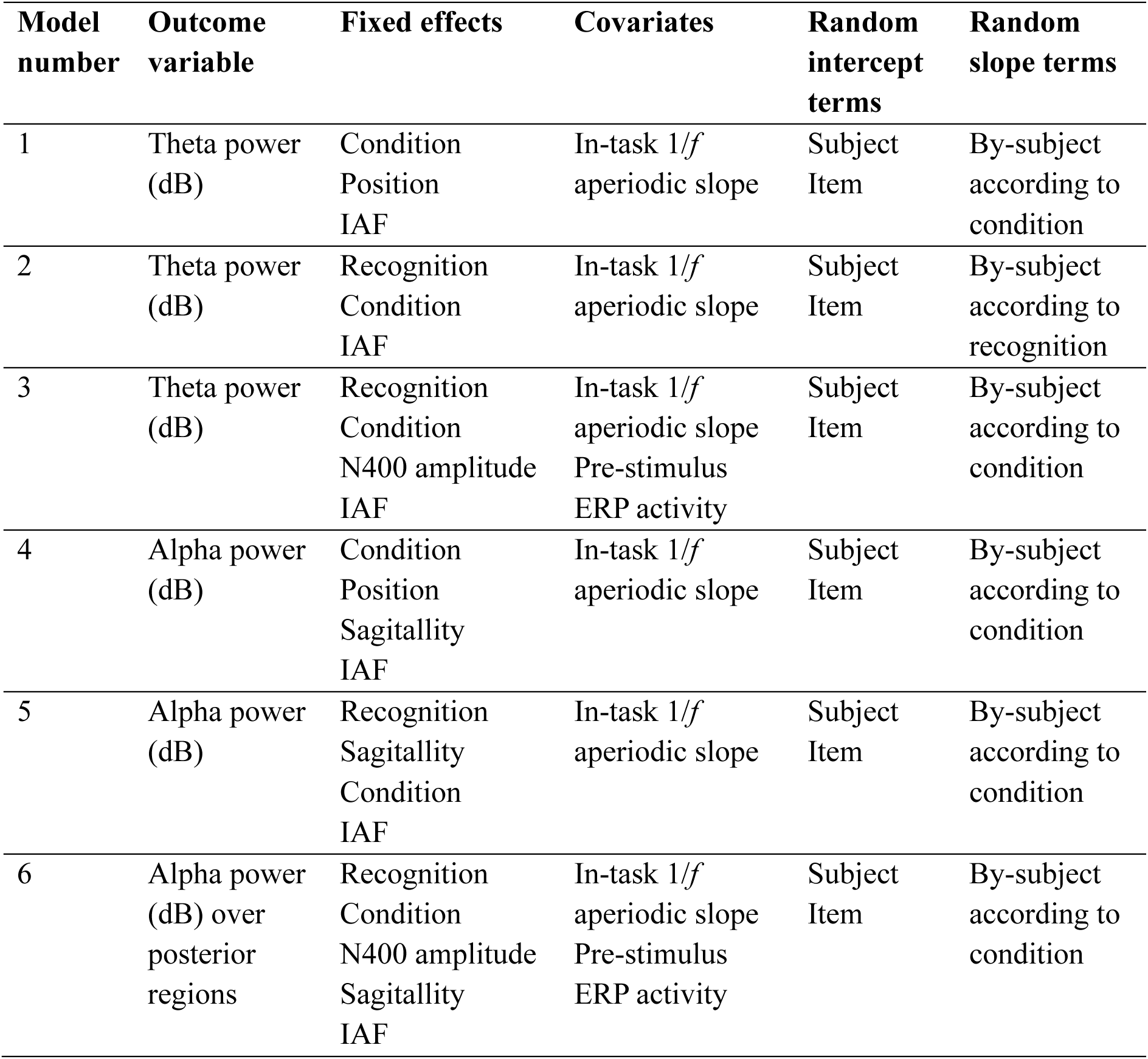
Linear mixed-effects model structures, including fixed and random effects. The levels of each variable include **condition** (predictable/unpredictable), **position** (A/B/C/D), **recognition** (hit/miss), and **sagitallity** (anterior/central/posterior).

| Model number | Outcome variable | Fixed effects | Covariates | Random intercept terms | Random slope terms |
| --- | --- | --- | --- | --- | --- |
| 1 | Theta power (dB) | Condition<br>Position<br>IAF | In-task 1/*f*<br>aperiodic slope | Subject<br>Item | By-subject according to condition |
| 2 | Theta power (dB) | Recognition<br>Condition<br>IAF | In-task 1/*f*<br>aperiodic slope | Subject<br>Item | By-subject according to recognition |
| 3 | Theta power (dB) | Recognition<br>Condition<br>N400 amplitude<br>IAF | In-task 1/*f*<br>aperiodic slope<br>Pre-stimulus<br>ERP activity | Subject<br>Item | By-subject according to condition |
| 4 | Alpha power (dB) | Condition<br>Position<br>Sagittality<br>IAF | In-task 1/*f*<br>aperiodic slope | Subject<br>Item | By-subject according to condition |
| 5 | Alpha power (dB) | Recognition<br>Sagittality<br>Condition<br>IAF | In-task 1/*f*<br>aperiodic slope | Subject<br>Item | By-subject according to condition |
| 6 | Alpha power (dB) over posterior regions | Recognition<br>Condition<br>N400 amplitude<br>Sagittality<br>IAF | In-task 1/*f*<br>aperiodic slope<br>Pre-stimulus<br>ERP activity | Subject<br>Item | By-subject according to condition |

## Results

Outputs from the statistical models are presented in Appendix A, and the data analysis scripts for the present reanalysis can be freely accessed here https://osf.io/zvpku/.

### Behavioural results from the memory recognition test

The number of hits (‘old’ responses to old images seen during the learning phase) and the number of misses (‘new’ responses to old images) were counted for each subject, condition, and position. Hit and miss rates were also calculated per participant and condition by dividing the number of hits and misses by the number of old items in each condition for each subject, to account for item imbalances between conditions (see Figure 2). Note that while hit rates alone are at or below chance, average d’ scores across image types (which are computed based on both old and new stimuli) were above chance in the prior study (greater than 0; Jano, Chatburn, et al., 2024). This may suggest that participants specifically experienced difficulty recognising old items, but not at rejecting new items.

**Figure 2.**
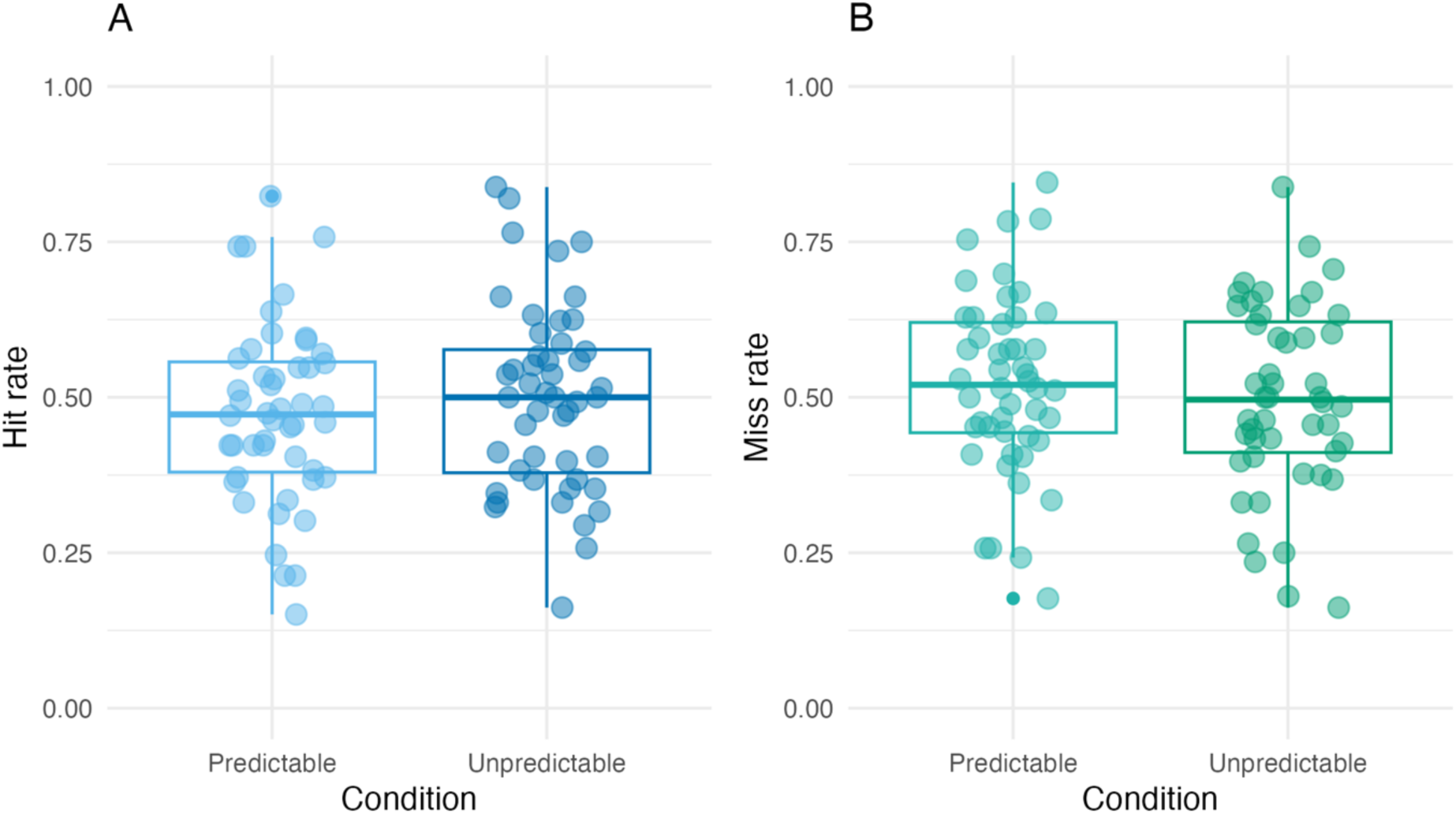
Boxplots depicting **(A)** hit rates and **(B)** miss rates between predictable and unpredictable image conditions. Data was extracted from performance during the recognition test. Dots reflect individual participants, centre lines within the boxes depict median values, with lines below and above reflecting the first and third quartiles, respectively.

### Visualising oscillatory and ERP patterns

Time frequency plots demonstrating raw data visualisations of oscillatory activity are presented in Figure 3 and Figure 4, along with the topography plots for theta and alpha power. Broadly, these plots demonstrate immediate power increases in both the theta and alpha oscillatory bands, followed by a reduction in power at the approximate 200ms to 250ms mark. While visual inspection of the heat maps shows differences between the conditions (predictable/unpredictable) and between the memory outcomes (hit/miss) in the alpha and theta bands, the most prominent differences appear in the beta frequency range (displayed in the difference plots). Grand average ERP plots according to subsequent memory performance do not reveal discernible differences in N400 amplitude to subsequently remembered versus forgotten images (see Figure 5). Note that this interaction alone was not a focus of the present study and was thus not examined in statistical analyses. For grand average ERP plots of the N400 across image conditions, the reader is directed to the original manuscript (Jano, Chatburn, et al., 2024).

**Figure 3.**
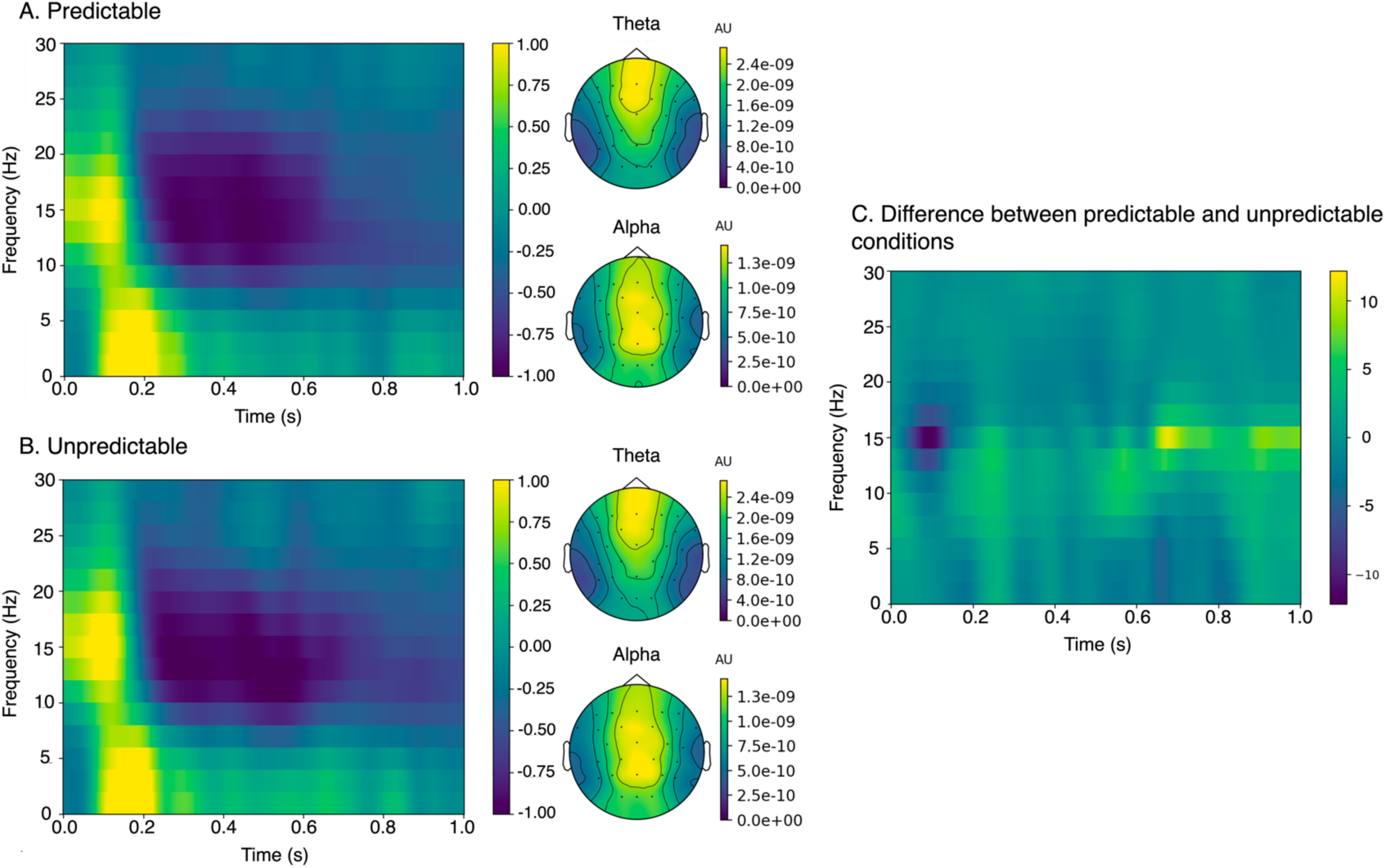
Time-frequency heat maps (displaying data averaged over all channels across the duration of image presentation; 0-1000ms) and topography plots according to **(A)** predictable and **(B)** unpredictable conditions. For heat maps, data across the whole epoch was divided by the data in the baseline window (−350 to -150ms pre-stimulus onset) and a decibel conversion was applied. Note that for statistical models, data in the 200-1000ms window post-image onset was used. For topography plots showing theta and alpha power, no baseline correction was applied. **(C)** Difference plot (predictable – unpredictable), showing z-scored, baseline corrected data (according to a -350 to -150ms pre-stimulus onset window).

**Figure 4.**
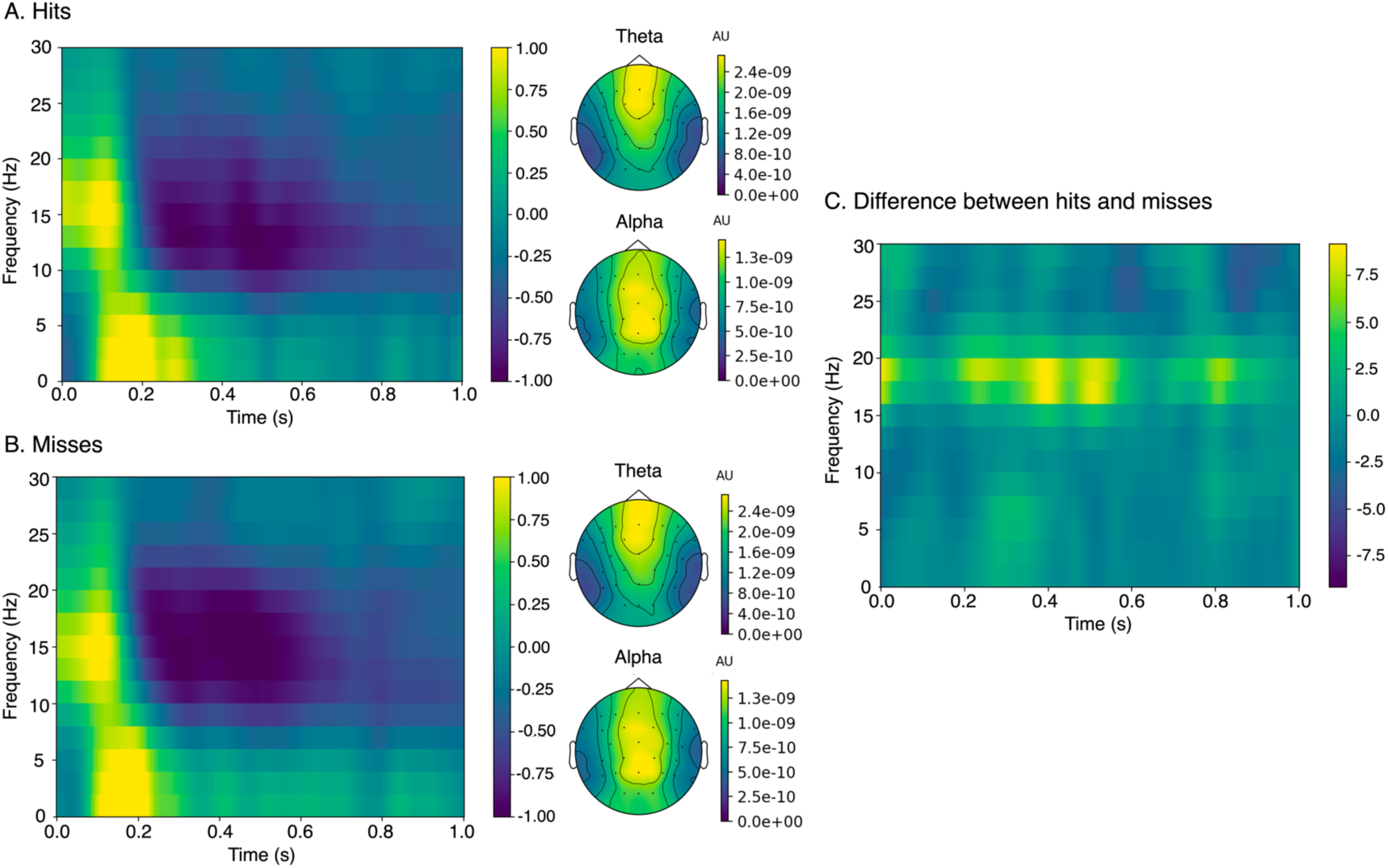
Time-frequency heat maps (displaying data averaged over all channels across the duration of image presentation; 0-1000ms) and topography plots according to **(A)** memory hits and **(B)** memory misses. For heat maps, data across the whole epoch was divided by the data in the baseline window (−350 to -150ms pre-stimulus onset) and a decibel conversion was applied. Note that for statistical models, data in the 200-1000ms window post-image onset was used. For topography plots showing theta and alpha power, no baseline correction was applied **(C)** Difference plot (hits – misses), showing z-scored, baseline corrected data (according to a -350 to -150ms pre-stimulus onset window).

**Figure 5.**
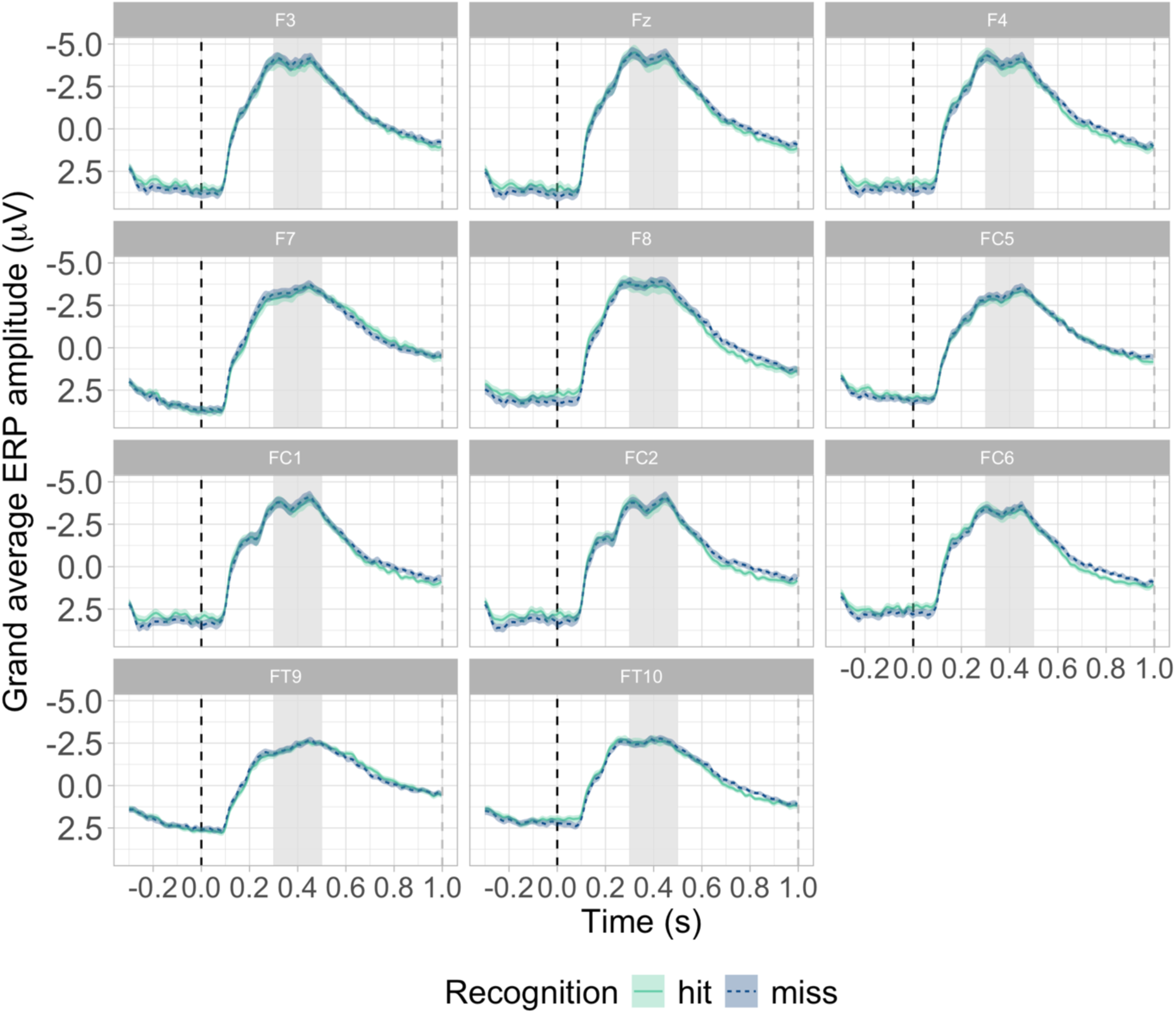
Grand average ERP amplitude at frontal channels according to time and subsequent memory performance. The black dotted line depicts image onset, while grey shading shows the N400 window. Solid turquoise lines reflect memory hits while dotted dark blue lines depict memory misses. Turquoise and dark blue shading represents the standard errors for the recognition types.

### Linear mixed model results of primary analyses

It was hypothesised that theta power during learning would be greatest for images in the unpredictable condition, as compared to images in the predictable condition (Hypothesis a). This was addressed via Model 1, which did not detect a significant main effect of *image condition* on theta power (χ^2^(1) = 0.98, *p* = .322). The interaction between *image condition, position* and *IAF* was also non-significant (χ^2^(3) = 5.09, *p* = .166), and as such, the hypothesis was unsupported.

The second hypothesis (Hypothesis b) predicted that alpha power during learning would differ between images in the predictable and unpredictable sequences (investigated via Model 4). While the main effect of *condition* on alpha power across all sagittal regions was non-significant (χ^2^(1) = 0.01, *p* = .925), the interaction between *condition, position* and *IAF* on alpha power was significant (χ^2^(3) = 8.42, *p* = .038). At ‘A’ images, higher IAFs were associated with greater alpha power across the predictable and unpredictable conditions. However, in the predictable condition, this relationship reversed at the ‘D’ image, with high IAFs being associated with decreases in alpha power (see Figure 6). As such, the hypothesis was partially supported, with the effect of image condition on alpha power depending on IAF and image position.

**Figure 6.**
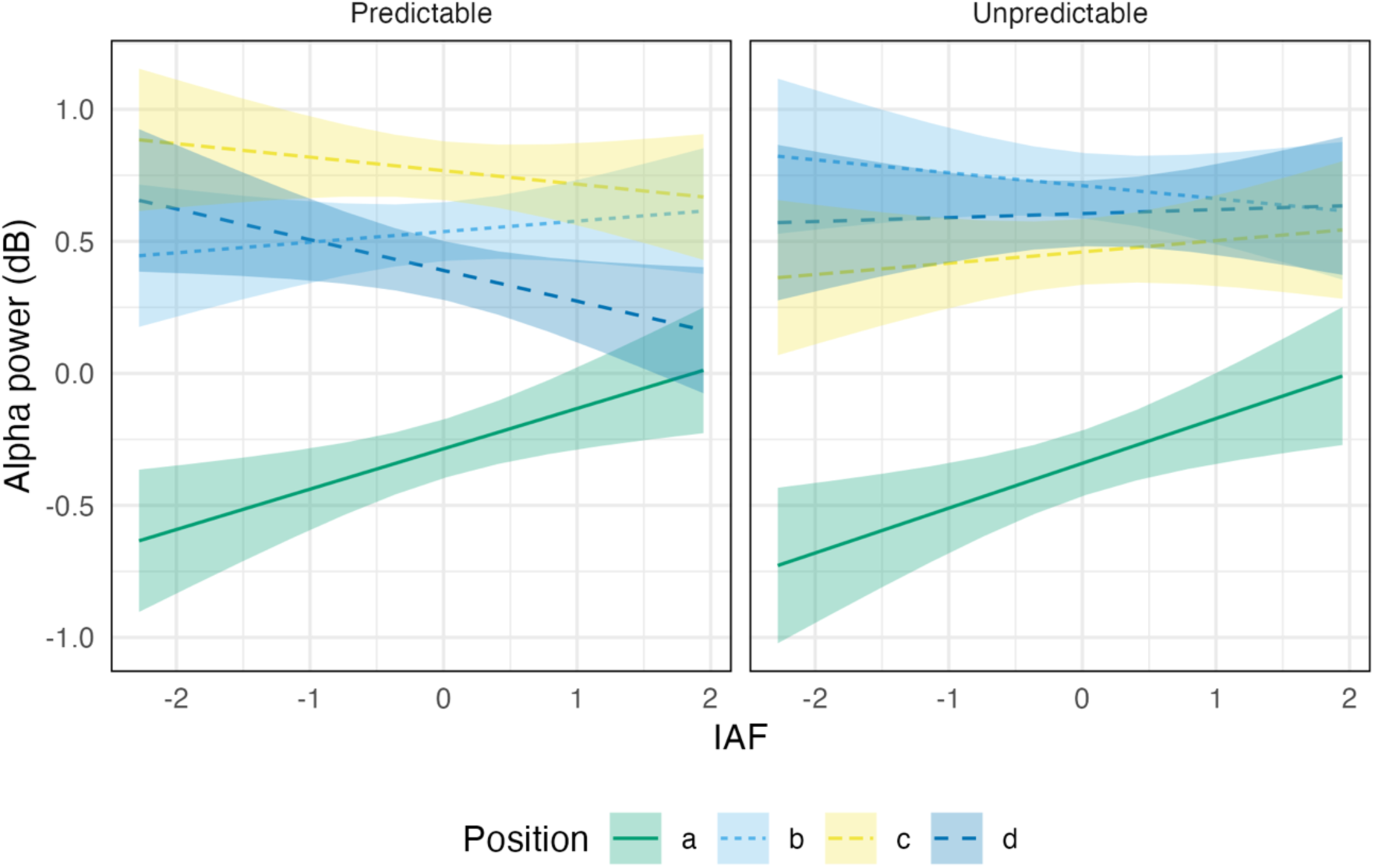
Linear mixed-effects model results of the relationship between alpha oscillatory power (across all sagittal regions), image position, image condition, and individual alpha frequency (IAF). Plots are faceted according to condition, whilst green, light blue, yellow and dark blue colours correspond to each image position (A, B, C, D, respectively). Shading represents 83% confidence intervals.

It was predicted that theta power during learning would be greatest for subsequently remembered items (memory hits) than subsequently forgotten items (memory misses; Hypothesis c). This was analysed via Model 2, which, in contrast to hypothesised effects, failed to detect a significant main effect of *memory recognition* on theta power (χ^2^(1) = 1.92, *p* = .166). The interaction between *recognition, IAF* and *condition* was also non-significant (χ^2^(1) = 0.04, *p* = .835), suggesting that theta power was not related to subsequent memory.

It was additionally predicted that alpha power during learning would be associated with memory recognition outcomes (Hypothesis d). This prediction was tested via model 5, which, in support of the hypothesis, revealed a significant main effect of *memory recognition* on alpha power across all sagittal regions (χ^2^(1) = 4.43, *p* = .035). Alpha desynchronisation was strongest for subsequently forgotten images (memory misses) as compared to subsequently remembered images (memory hits). However, the interaction between *recognition, sagitallity, condition* and *IAF* was non-significant (χ^2^(2) = 0.11, *p* = .948). Consequently, the relationship between alpha power and memory encoding may not depend on IAF or N400 amplitudes.

It was also hypothesised that the relationship between the N400 and oscillatory activity during learning would be linked to subsequent memory outcomes at test (Hypothesis e). Model 3 was computed to test this hypothesis in relation to theta power, and, in support of the hypothesis, revealed a significant interaction between *memory recognition, IAF* and *N400 amplitude* on theta power (χ^2^(1) = 4.24, *p* = .039). For low IAF individuals, subsequently forgotten items (memory misses) were associated with increases in theta power as the N400 amplitude became larger (more negative). For subsequently remembered items (memory hits), this relationship was present albeit weaker. In contrast, for high IAF individuals, subsequently forgotten items (memory misses) were associated with decreased theta power as the N400 amplitude increased. Subsequently remembered items (memory hits) were associated with a weak relationship between theta and the N400 amplitude (see Figure 7). However, the main effect of N400 amplitude on theta power was nonsignificant (χ^2^(1) = 2.07, *p* = .150). Therefore, while N400 amplitudes alone were unrelated to changes in theta power, the N400, theta power and IAF interacted to influence later remembering.

**Figure 7.**
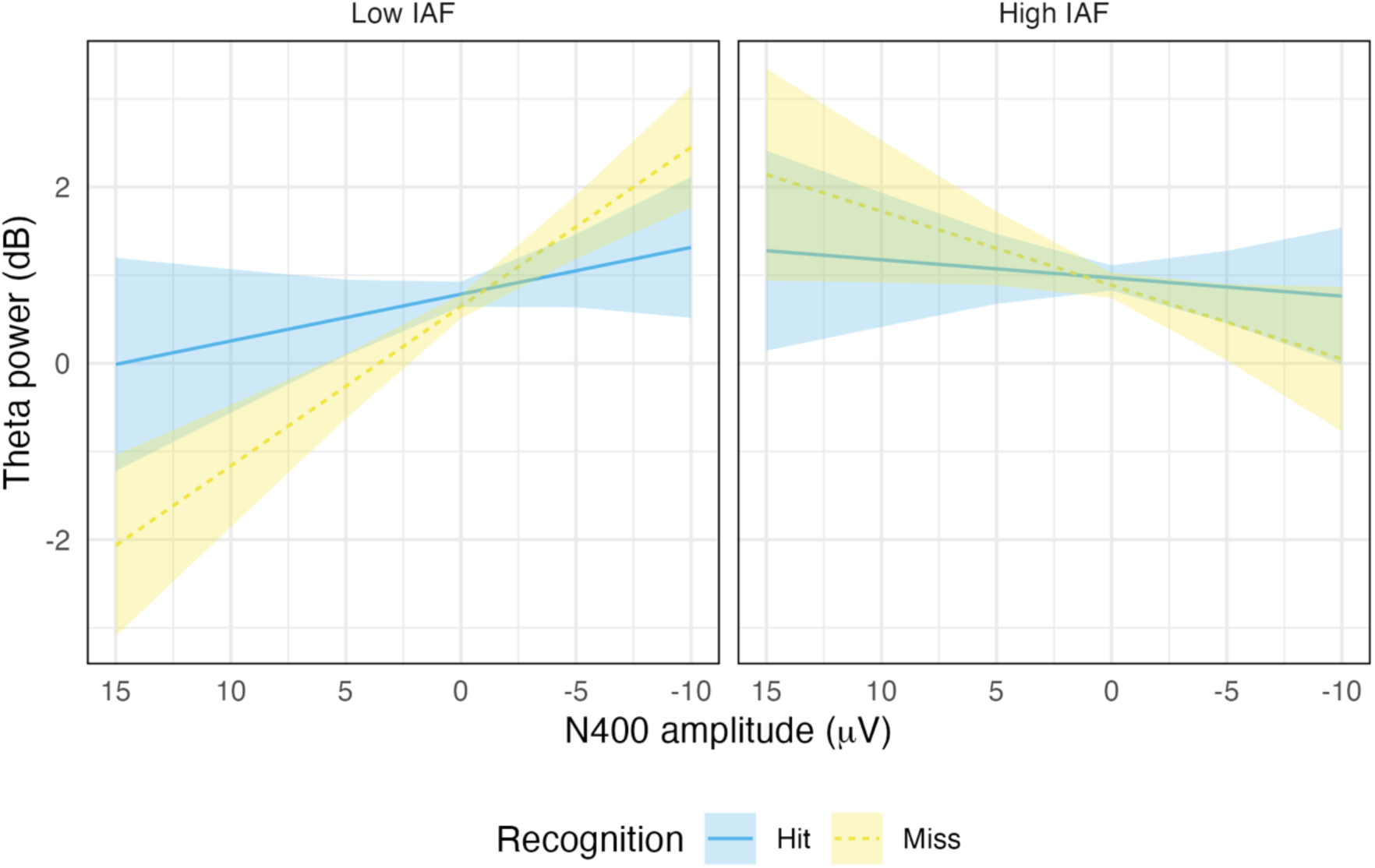
Linear mixed-effects model results depicting the relationship between theta power, N400 amplitude, memory recognition, and individual alpha frequency (IAF). Blue lines reflect subsequently remembered items (memory hits) whilst yellow dotted lines represent subsequently forgotten items (memory misses). The plot is faceted into low and high IAF, with this dichotomisation for visualisation purposes only.

Finally, Model 6 tested the hypothesis concerning oscillatory and N400 subsequent memory effects (Hypothesis e) in relation to alpha power. Contrary to hypothesised effects, the model did not detect a significant interaction between *memory recognition, N400 amplitude, IAF, condition* and *sagitallity* on alpha power (χ^2^(2) = 1.04, *p* = .594). The main effect of N400 amplitude on alpha power was also nonsignificant (χ^2^(1) = 0.13, *p* = .720). As such, alpha power did not interact with N400 amplitude to influence subsequent memory, and N400 amplitude alone was not associated with changes in alpha power.

### Exploratory correlation analysis

As primary models revealed that memory hits and misses were supported by differential patterns of neural activity between high and low IAF individuals, we sought to determine whether such patterns might be related to increased or decreased hit rates. As such, a Pearson correlation was conducted to explore the relationship between IAF and hit rate, after Shapiro-Wilk tests revealed that the variables were normally distributed. The Pearson correlation revealed a significant but weak relationship between the two variables (*r*(44)= 0.36, *p* = .014), such that increases in IAF were associated with increasing hit rates (see Figure 8).

**Figure 8.**
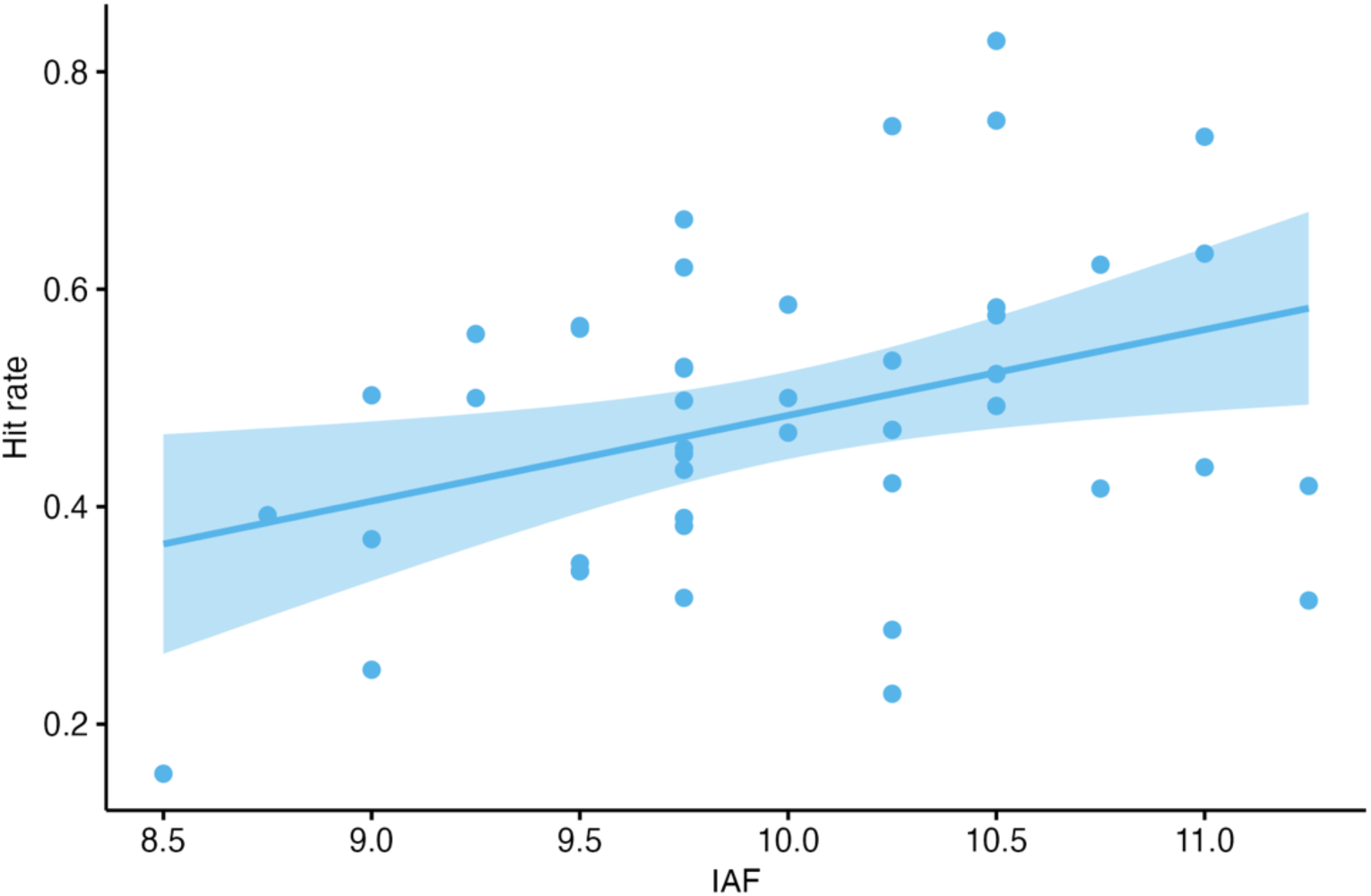
The correlation between individual alpha frequency (IAF) and hit rate (across all image positions and conditions). Dots represent individual data points (N = 46).

## Discussion

The present investigation reanalysed data from a previous study (Jano, Chatburn, et al., 2024) to further elucidate the relationship between prediction and memory. Participants’ brain activity was recorded using EEG whilst they viewed predictable and unpredictable sequences of outdoor scene images. Electrophysiological activity during learning was then compared for images subsequently recognised and forgotten. Contrary to expectations, theta power did not differ between images in unpredictable and predictable conditions. However, theta power interacted with IAF and the N400 component to influence later remembering. Based on the strong relationship between the N400 and item expectancy (DeLong et al., 2005; for a review see Kutas & Federmeier, 2011), the present findings could imply that intrinsic neural variability modulates the degree to which prediction errors drive memory encoding. Additionally, alpha desynchronisation was strongest for items later forgotten versus remembered. This reiterates the functional relevance of alpha power for learning and information uptake. Ultimately, the results emphasise the importance of ERPs and oscillatory rhythms for memory formation, whilst suggesting that the effect of prediction on memory differs at the inter-individual level.

### Theta power alone does not relate to subsequent memory

Oscillatory theta power has previously been linked to successful memory (Klimesch, 1999; Klimesch et al., 1997; Sederberg et al., 2003) and to prediction error processing (Cavanagh et al., 2010, 2012), suggesting that theta is useful in gauging the memory updating processes related to prediction. Given also that prediction and encoding may be inversely related (Sherman et al., 2022; Sherman & Turk-Browne, 2020), it was hypothesised that theta power would be stronger for images in the unpredictable sequences as compared to the predictable sequences, reflecting heightened memory encoding in the absence of a strong prediction. While the expected trend was somewhat evident in the present time-frequency plots, this between-condition difference in theta synchronisation did not reach significance. At a first glance, this suggests that predictions do not influence theta synchronisation (and by extension, memory encoding). However, as discussed in Jano, Chatburn, and colleagues (2024), it is also likely that the categorical manipulation of predictability in the present experiment was too superficial to capture the nuanced changes in stimulus expectancy present in the dataset. Given also that each image was unique (despite belonging to either a predictable or unpredictable categorical sequence), predictability at the item level may not have been measured. This may explain the failure of the present study to detect a condition-specific effect relating to theta power, as memory encoding might have occurred in both conditions. However, the relationship between theta, prediction, and memory is more clearly illuminated via the effects relating to subsequent memory.

It was predicted that increases in theta power during learning would be associated with successful remembering at test. However, in contrast to previous literature linking theta synchronisation to enhanced subsequent memory (e.g. Klimesch, Doppelmayr, et al., 1996; Long et al., 2014; Sederberg et al., 2003), the present analysis did not reveal a relationship between theta power and memory performance. Whilst theta power during learning did not influence later remembering, this finding may relate to the level of processing required by the present task. Subsequent memory effects of oscillatory patterns are largely variable and task-dependent, as the type of paradigm employed may give rise to varying depths of processing, consequently influencing the extent of memory encoding (Hanslmayr et al., 2009; Hanslmayr & Staudigl, 2014). The present study employed an incidental memory encoding paradigm with realistic, complex outdoor scene images, as we sought to examine the consequences of prediction for memory as predictions naturally develop. This is in contrast to prior research reporting theta subsequent memory effects, which often uses more simplistic stimuli (e.g., word lists or pictures of objects), informs participants that their memory will later be tested, instructs participants to remember the content, or requires that they answer a question about each item during learning (e.g., Klimesch, Doppelmayr, et al., 1996; Long et al., 2014; White et al., 2013). Such procedures may encourage memory encoding to a greater extent than the present paradigm (although may be limited in their ability to gauge more naturalistic forms of learning and prediction).

This is furthered by the low hit rates observed in the behavioural data (at approximately 0.50), suggesting that overall encoding in the current study may have been weak. However, it is important to note that the original study by Sherman & Turk-Browne (2020), upon which the present design was based, obtained similar average hit rate values (ranging from 0.38 to 0.61). Furthermore, while lower hit rates might be problematic for research primarily interested in behavioural memory performance, the focus of the present investigation was on the electrophysiological patterns that relate to later remembering and forgetting. As such, we do not view the lower hit rates as a limitation of the paradigm per se, but instead as a novel contribution to the existing literature, highlighting the limits of memory formation under more complex and dynamic circumstances. Additionally, as d’ scores in the previous study were above chance (Jano, Chatburn, et al., 2024), the lower hit rate implies a specific encoding decrement as opposed to overall impaired memory performance. Consequently, future research could vary the level of stimulus complexity to determine the bounds of memory formation in more naturalistic scenarios. Moreover, since the learning in the current study resembles learning in an everyday context, where individuals are not explicitly told to remember or predict information, day-to-day memory performance may be weaker compared to that observed in experimental settings. Importantly, these results alone do not speak directly to effects relating to prediction, particularly given the potential for varied stimulus predictability levels within both the predictable and unpredictable image conditions. This further underscores the importance of considering the N400, which may capture the extent of predictive processing more directly due to its ties to prediction error.

### IAF may modulate the relationship between prediction and memory encoding

The current study is the first to our awareness to directly compare the relationship between oscillatory and N400 patterns during a prediction task with subsequent memory performance and intrinsic neural factors. Interestingly, when the N400 component was incorporated into models of theta power, IAF and subsequent memory, a relationship between the variables was observed. For individuals with a low IAF, increases in N400 amplitude (becoming more negative) were associated with increases in theta power, with this relationship being strongest for subsequently forgotten items (memory misses) versus subsequently remembered items (memory hits). Prior research links the N400 to prediction error (e.g., Bornkessel-Schlesewsky & Schlesewsky, 2019; Eddine et al., 2024; Rabovsky & McRae, 2014). Therefore, the present results could indicate that for low IAF individuals, optimal remembering is supported by a weak positive relationship between the magnitude of prediction error and theta power during learning. This implies that for such individuals, ‘too much’ encoding in response to a prediction error, or ‘too little’ encoding in the absence of a prediction error is maladaptive for later remembering. Conversely, for high IAF individuals, successful remembering was supported by the absence of a relationship between theta power and the N400 during learning. Moreover, subsequently forgotten stimuli were associated with a stronger *negative* relationship between theta power and the magnitude of the N400, suggesting that ‘too little’ encoding in the face of a prediction error and ‘too much’ encoding in response to predictable information is detrimental. This could point more generally to an ‘optimal’ level of prediction error-related memory uptake, which appears to differ between individuals. Here, over-encoding might lead to interference in memory, while under-encoding might prevent the information from entering memory. However, the underlying reason for these differential effects across varying levels of predictability and IAF remains uncertain, rendering it challenging to draw confident conclusions. Nevertheless, the effects associated with successful memory might be illuminated by a consideration of how those with high and low IAFs differ with regard to cognitive processing.

Higher IAFs are associated with better general intelligence (Grandy et al., 2013), enhanced memory (Klimesch, 1997; Klimesch et al., 1990, 1993), and faster perceptual processing windows (Samaha & Postle, 2015). Although IAF has particularly been linked to the speed of memory retrieval, it has also exhibited differences during memory encoding between good and bad performers, suggesting that those with a high IAF have a heightened memory encoding ability (Klimesch, 1997; Klimesch et al., 1993). Consequently, the observation that successful memory was not supported by the relationship between theta and the N400 for those with a high IAF may imply that such individuals rely less on prediction error signals to inform memory updating than their low IAF counterparts. This is compatible with our previous findings, where N400 amplitudes did not relate to memory for those with a high IAF, prompting the suggestion that such individuals adopt a more widespread encoding strategy (Jano, Chatburn, et al., 2024). Moreover, the present results suggest that this more widespread “strategy” can have benefits for memory performance, as IAF and hit rate were (allbeit weakly) positively correlated. To build upon this suggestion, if high IAF individuals are better at encoding information more generally, prediction errors may not contribute over and above to the memory encoding that is already occurring (hence the weak relationship between theta and the N400 amplitude observed for memory hits). While the current analysis cannot speak directly to how these disparate memory strategies may manifest, it may be a useful first step in explaining why at times, memory is enhanced in cases of low prediction error (i.e., for predictable stimuli; e.g., Gronau & Shachar, 2015; Turan et al., 2023), but also for unexpected (i.e., prediction error-eliciting) information (Federmeier et al., 2007; Rouhani et al., 2018).

Although the relationship between prediction and memory appears to be modulated by IAF, it is important to emphasise that the present study could not explicitly relate the amplitude of the N400 to fluctuations in image predictability due to the limits of the categorical predictability manipulation. As such, the extent to which the N400 reflects a prediction error signal in this context is unclear, although the generalisability of this component to non-linguistic settings is highly promising (for discussion see Eddine et al., 2024; Kutas & Federmeier, 2011). Similarly, as theta power alone did not relate to subsequent memory, it is possible that such activity was not reflective of memory encoding in the present study. As visual inspection of the time frequency heat maps reveals strong beta power differences between subsequently remembered and forgotten images, activity in this frequency band might have related more strongly to memory encoding. Consequently, an analysis of the beta frequency band, and it’s relationship to prediction and memory, may be an interesting endeavour for future research. However, returning to a discussion of the theta rhythm, it is likely that theta subsequent memory effects were generally smaller than in previous studies and as such, only detectable once other sources of variability (such as N400 amplitude) were considered. Similarly, grand average ERP plots did not show discernible differences in N400 amplitude according to subsequently remembered and forgotten stimuli. This could suggest that the effect of the N400 on memory is also dependent on other factors; a proposal that is supported by the variable relationship observed here according to theta power and IAF. Nevertheless, these findings imply that the association between theta power and the N400 is functionally significant for later remembering, although future research may benefit from more fine-grained predictability measurements. Ultimately, the current results emphasise the more intricate relationship between prediction and memory; while prediction errors could promote memory formation, the mixed findings according to IAF also cannot rule out the potential for encoding to occur independently of, or over and above, the level of prediction error. This suggestion is further reiterated by the results pertaining to alpha power.

### The effect of alpha power on memory encoding does not relate to the N400

To provide further insight into oscillatory changes during predictive processing, alpha activity during the learning phase was measured. The present results revealed that IAF and alpha power were positively related at the ‘A’ images (regardless of image condition), and negatively related at predictable ‘D’ images. It is possible that this pattern arises from differences in memory encoding during naturalistic sequence processing, potentially stemming from changes in predictability. At the beginning of the sequence, it was likely difficult to determine whether the sequence would be predictable or unpredictable. However, this would be clear by the end of the sequence, where the observed items either matched or mismatched previously learnt regularities. The decreased alpha power observed following predictable ‘D’ images for high IAF individuals may suggest that alpha activity adapted according to changes in stimulus relevance and predictability. Alpha desynchronisation has been linked to successful memory formation (Hanslmayr et al., 2009; Klimesch, Schimke, et al., 1996; Sederberg et al., 2003), and as such, high IAF individuals may have been more likely to encode predictable ‘D’ images into memory than low IAF individuals. This is compatible with the above suggestion that such individuals have a more widespread propensity for memory encoding. Alternatively, *increased* alpha synchronisation could promote memory encoding because it inhibits the retrieval of task-irrelevant memory representations (Klimesch et al., 2007). With this in mind, those with a high IAF, upon encountering a predictable stimulus, may have directed their encoding resources away from the redundant information. However, while this result points toward a potential optimisation of processing according to stimulus predictability and intrinsic neural differences, the effects relating to alpha and subsequent memory can permit more direct conclusions regarding alpha’s relevance for memory encoding.

Greater alpha desynchronisation (i.e., reduced alpha power) during learning was associated more strongly with forgetting versus successful remembering. This is in line with previous research (Meeuwissen et al., 2011), and with alpha’s proposed relation to inhibitory processing (Klimesch et al., 2007), suggesting that increases in alpha power during learning positively influence memory formation. However, alpha subsequent memory effects in the present study were not related to image condition or to the amplitude of the N400 component, providing further support for the notion that prediction error may not be a necessary prerequisite for memory formation. Given also that N400 amplitudes were not correlated with alpha power, the two components appear to reflect differing mechanisms. Taken together, these findings have implications for predictive processing theories, which conceptualise prediction as a general coding strategy (Huang & Rao, 2011). Prediction errors are proposed to signal a need for model updating and may be minimized through changes in neuronal connections (i.e., synaptic plasticity; Barron et al., 2020; den Ouden et al., 2012; Friston, 2005). These neuronal alterations purportedly underlie memory encoding, suggesting that prediction errors drive learning (Friston, 2005; for a review of synaptic plasticity and memory see Martin et al., 2000). However, while prediction may be a general cognitive mechanism, the present results question the role of prediction errors as the primary learning signals in the brain. As it is difficult to determine the extent to which the present categorical predictability manipulation captures subtle changes in predictability, a design that adjusts stimulus probabilities at a more fine-grained level could provide stronger insights into how changes in oscillatory and ERP activity relate to prediction and subsequent memory.

## Conclusion

The current study emphasises the dynamic and inter-individual nature of the relationship between prediction and memory. Although likely interconnected, memory encoding may not depend completely on the magnitude of prediction error, adding a layer of complexity to the potential trade-off between prediction and memory uptake (Hubbard et al., 2019; Sherman et al., 2022; Sherman & Turk-Browne, 2020). Most notably, whilst low IAF individuals appear more inclined to update their predictive memory representations according to prediction errors, individuals with a high IAF could rely on a more general memory updating strategy that is independent of prediction error, or that contributes to memory encoding over and above the model updating resulting from prediction errors. Although this may have significant implications for predictive coding schemes, it also presents intriguing opportunities for future research, which should further examine individual neural variability in the context of prediction and memory. Finally, the present reanalysis provides evidence for an inherent link between oscillatory activity, IAF, N400 amplitude, and subsequent memory, whilst also suggesting that oscillations in the alpha and theta bands reflect somewhat distinct processes to the N400 ERP.

## Appendix A: Statistical model outputs

### Model 1: theta_db_mean ∼ condition * position * cog_scale + intask_slope_scale + (1+condition|subjnr) + (1|item_no), data= theta_sum

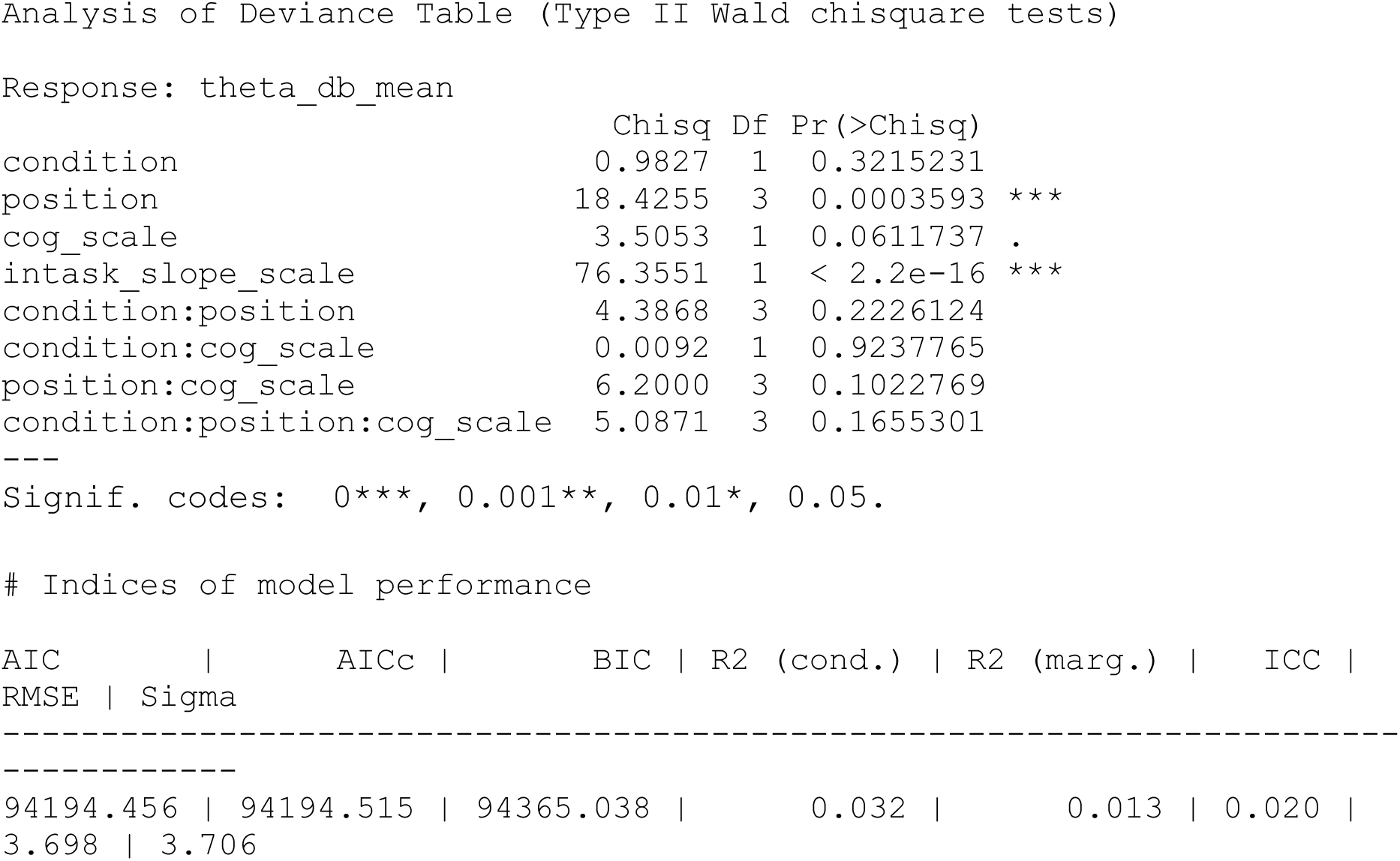

### Model 2: theta_db_mean ∼ recognition * cog_scale * condition + intask_slope_scale + (1+recognition|subjnr) + (1|item_no), data= theta_sum

|  | Chisq | Df | Pr(>Chisq) |
| --- | --- | --- | --- |
| recognition | 1.9161 | 1 | 0.16629 |
| cog_scale | 4.0538 | 1 | 0.04407 * |
| condition | 0.5320 | 1 | 0.46579 |
| intask_slope_scale | 81.6596 | 1 | < 2e-16 *** |
| recognition:cog_scale | 0.0719 | 1 | 0.78864 |
| recognition:condition | 0.1046 | 1 | 0.74640 |
| cog_scale:condition | 0.0274 | 1 | 0.86851 |
| recognition:cog_scale:condition | 0.0432 | 1 | 0.83535 |

Indices of model performance
| AIC | AICc | BIC | R2 (cond.) | R2 (marg.) | ICC |
| --- | --- | --- | --- | --- | --- |
| RMSE | Sigma |  |  |  |  |
| --- | --- | --- | --- | --- | --- |
| 93182.985 | 93183.010 | 93291.391 | 0.030 | 0.011 | 0.019 |
| 3.701 | 3.708 |  |  |  |  |

Analysis of Deviance Table (Type II Wald chisquare tests)
Response: $\theta_{db\_mean}$

### Model 3: theta_db_mean ∼ recognition * N400_scale * cog_scale * condition + intask_slope_scale + prestim_scale + (1+condition|subjnr) + (1|item_no), data= theta_sum

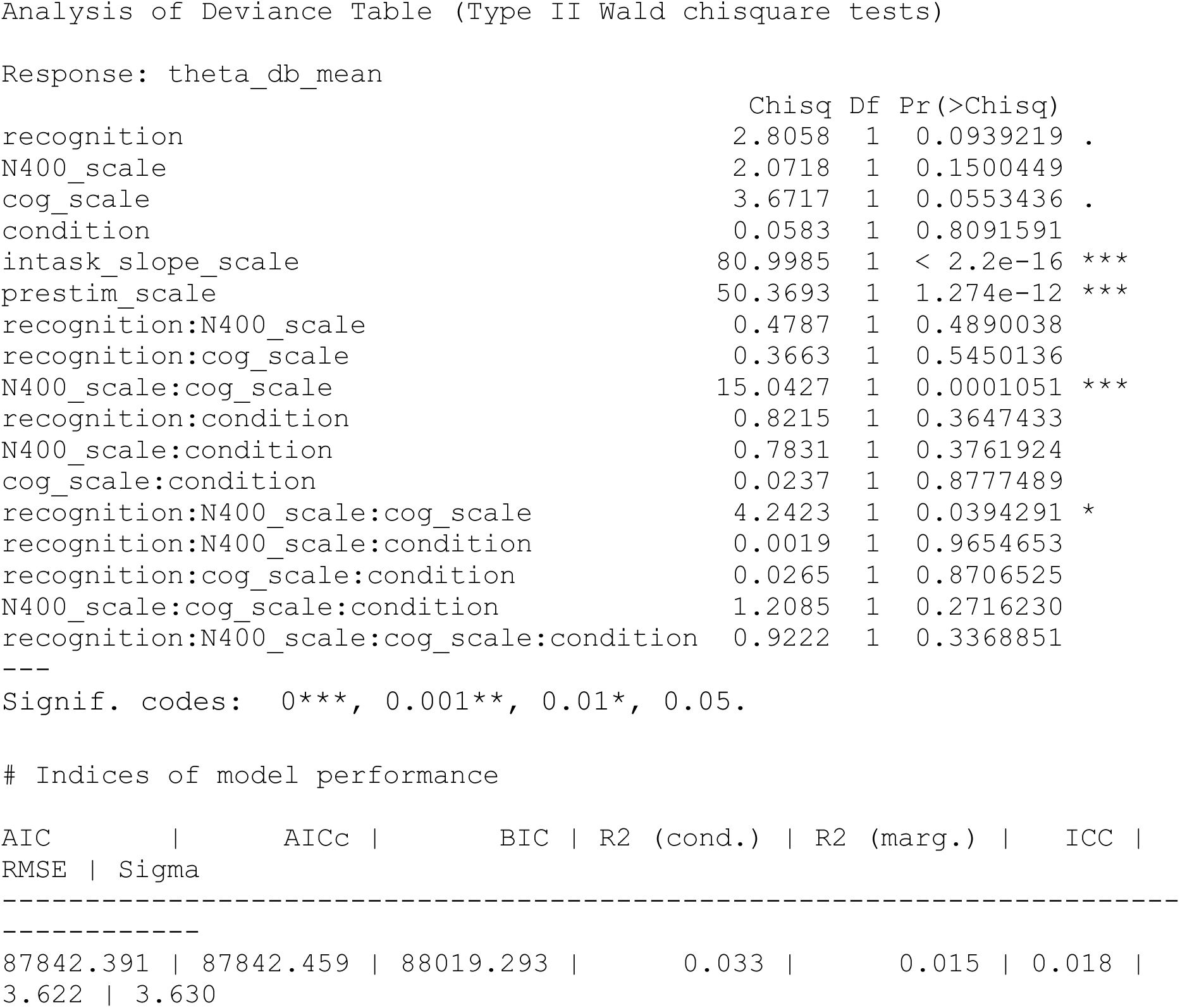

### Model 4: alpha_db_mean ∼ position * condition * cog_scale * sag. + intask_slope_scale + (1+condition|subjnr) + (1|item_no), data= region_sum

|  | Chisq | Df | Pr(>Chisq) |  |
| --- | --- | --- | --- | --- |
| position | 818.8705 | 3 | < 2.2e-16 | *** |
| condition | 0.0088 | 1 | 0.92513 |  |
| cog_scale | 0.1158 | 1 | 0.73366 |  |
| sag. | 22.5502 | 2 | 1.269e-05 | *** |
| intask_slope_scale | 3.3245 | 1 | 0.06825 | . |
| position:condition | 52.9300 | 3 | 1.897e-11 | *** |
| position:cog_scale | 38.7955 | 3 | 1.918e-08 | *** |
| condition:cog_scale | 0.5697 | 1 | 0.45039 |  |
| position:sag. | 150.0279 | 6 | < 2.2e-16 | *** |
| condition:sag. | 0.2828 | 2 | 0.86816 |  |
| cog_scale:sag. | 26.3612 | 2 | 1.887e-06 | *** |
| position:condition:cog_scale | 8.4154 | 3 | 0.03816 | * |
| position:condition:sag. | 8.7738 | 6 | 0.18670 |  |
| position:cog_scale:sag. | 15.2049 | 6 | 0.01872 | * |
| condition:cog_scale:sag. | 0.9283 | 2 | 0.62866 |  |
| position:condition:cog_scale:sag. | 1.1034 | 6 | 0.98139 |  |

Indices of model performance
| AIC | AICc | BIC | R2 (cond.) | R2 (marg.) | ICC |
| --- | --- | --- | --- | --- | --- |
| RMSE | Sigma |  |  |  |  |
| ----- |  |  |  |  |  |
| 2.635e+05 | 2.635e+05 | 2.640e+05 | 0.047 | 0.021 | 0.027 |
| 3.077 | 3.083 |  |  |  |  |

Analysis of Deviance Table (Type II Wald chisquare tests)
Response: $\alpha_{db\_mean}$

### Model 5: alpha_db_mean ∼ recognition * sag. * condition * cog_scale + intask_slope_scale + (1+condition|subjnr) + (1|item_no), data= region_sum

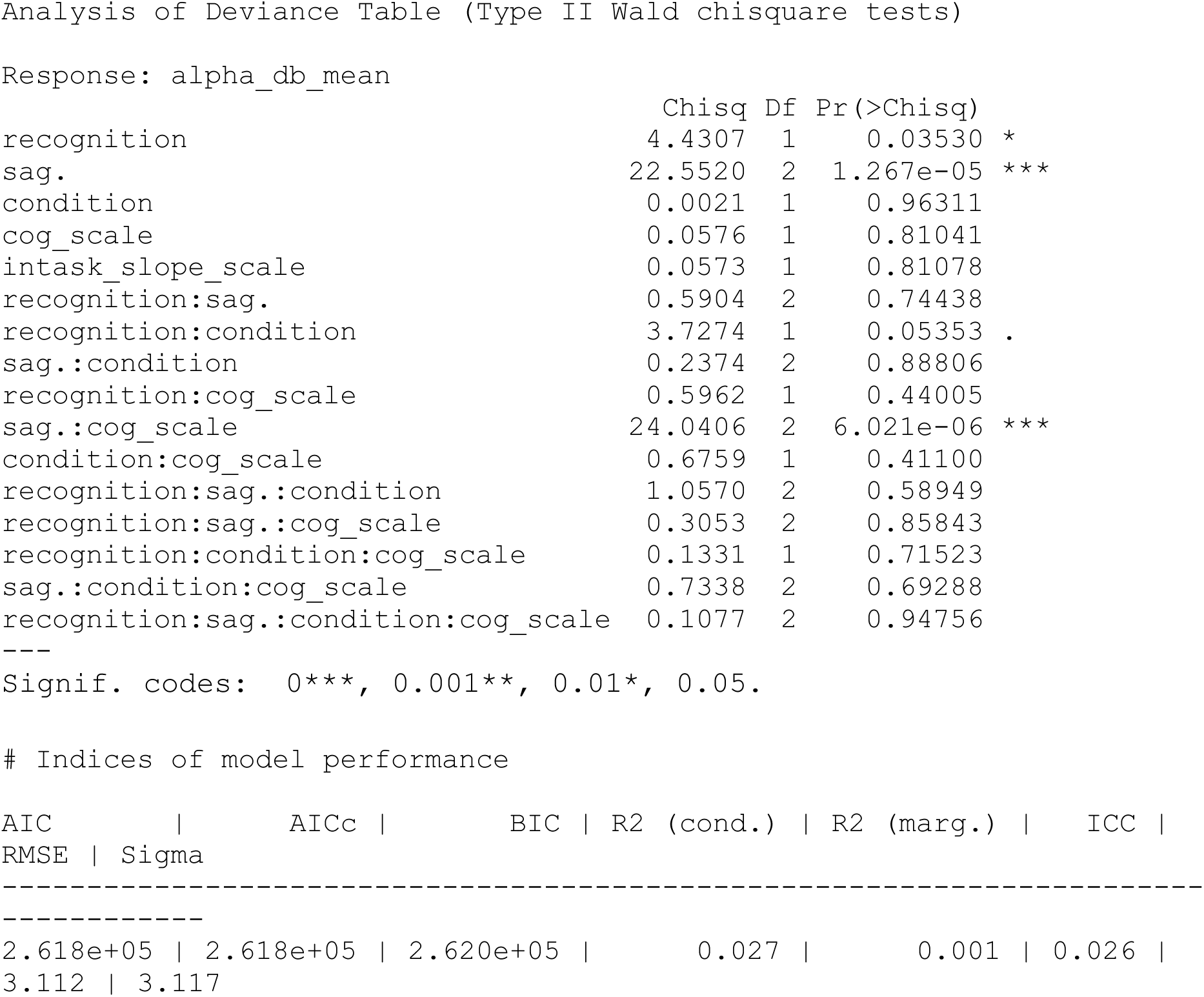

### Model 6: alpha_db_mean ∼ recognition * N400_scale * sag. * cog_scale * condition + prestim_scale + intask_slope_scale + (1+condition|subjnr) + (1|item_no), data= region_sum

|  | Chisq | Df | Pr(>Chisq) |
| --- | --- | --- | --- |
| recognition | 6.2839 | 1 | 0.01218 * |
| N400_scale | 0.1288 | 1 | 0.71966 |
| sag. | 20.7551 | 2 | 3.112e-05*** |
| cog_scale | 0.0161 | 1 | 0.89899 |
| condition | 0.0738 | 1 | 0.78589 |
| prestim_scale | 15.8839 | 1 | 6.735e-05*** |
| intask_slope_scale | 1.2628 | 1 | 0.26112 |
| recognition:N400_scale | 0.4087 | 1 | 0.52262 |
| recognition:sag. | 0.4092 | 2 | 0.81498 |
| N400_scale:sag. | 0.2021 | 2 | 0.90390 |
| recognition:cog_scale | 1.0000 | 1 | 0.31730 |
| N400_scale:cog_scale | 16.0917 | 1 | 6.035e-05*** |
| sag.:cog_scale | 20.7931 | 2 | 3.054e-05*** |
| recognition:condition | 0.7711 | 1 | 0.37988 |
| N400_scale:condition | 1.4623 | 1 | 0.22656 |
| sag.:condition | 0.2449 | 2 | 0.88474 |
| cog_scale:condition | 0.6889 | 1 | 0.40654 |
| recognition:N400_scale:sag. | 0.2311 | 2 | 0.89086 |
| recognition:N400_scale:cog_scale | 3.7515 | 1 | 0.05276 . |
| recognition:sag.:cog_scale | 0.3747 | 2 | 0.82916 |
| N400_scale:sag.:cog_scale | 3.1968 | 2 | 0.20222 |
| recognition:N400_scale:condition | 0.4468 | 1 | 0.50384 |
| recognition:sag.:condition | 1.1473 | 2 | 0.56346 |
| N400_scale:sag.:condition | 1.0961 | 2 | 0.57809 |
| recognition:cog_scale:condition | 0.0317 | 1 | 0.85879 |
| N400_scale:cog_scale:condition | 0.0734 | 1 | 0.78650 |
| sag.:cog_scale:condition | 0.3274 | 2 | 0.84902 |
| recognition:N400_scale:sag.:cog_scale | 1.5689 | 2 | 0.45637 |
| recognition:N400_scale:sag.:condition | 0.1864 | 2 | 0.91103 |
| recognition:N400_scale:cog_scale:condition | 0.6700 | 1 | 0.41305 |
| recognition:sag.:cog_scale:condition | 0.0277 | 2 | 0.98625 |
| N400_scale:sag.:cog_scale:condition | 1.1490 | 2 | 0.56297 |
| recognition:N400_scale:sag.:cog_scale:condition | 1.0418 | 2 | 0.59399 |

Indices of model performance
| AIC | AICc | BIC | R2 (cond.) | R2 (marg.) | ICC |
| --- | --- | --- | --- | --- | --- |
| RMSE | Sigma |  |  |  |  |
| 2.467e+05 | 2.467e+05 | 2.471e+05 | 0.027 | 0.002 | 0.024 |
| 3.045 | 3.051 |  |  |  |  |

## Reference

1. Alday, P. M. (2018). Philistine (v0.1) [Source code]. Available online at: https://github.com/palday/philistine/

2. Amoruso, L., Gelormini, C., Aboitiz, F., Alvarez González, Miguel, Manes, F., Cardona, J., & Ibanez, A. (2013). N400 ERPs for actions: Building meaning in context. Frontiers in Human Neuroscience, 7. https://www.frontiersin.org/articles/10.3389/fnhum.2013.00057

3. Baker, R., Dexter, M., Hardwicke, T. E., Goldstone, A., & Kourtzi, Z. (2014). Learning to predict: Exposure to temporal sequences facilitates prediction of future events. Vision Research, 99, 124–133. 10.1016/j.visres.2013.10.017

4. Bar, M. (2007). The proactive brain: Using analogies and associations to generate predictions. Trends in Cognitive Sciences, 11(7), 280–289. 10.1016/j.tics.2007.05.005

5. Bar, M. (2009). The proactive brain: Memory for predictions. Philosophical Transactions of the Royal Society B: Biological Sciences, 364(1521), 1235–1243. 10.1098/rstb.2008.0310

6. Barron, H. C., Auksztulewicz, R., & Friston, K. (2020). Prediction and memory: A predictive coding account. Progress in Neurobiology, 192, 101821. 10.1016/j.pneurobio.2020.101821

7. Bates, D., Kliegl, R., Vasishth, S., & Baayen, H. (2018). Parsimonious Mixed Models. *arXiv:1506.04967 [Stat]*. http://arxiv.org/abs/1506.04967

8. Bates, D., Mächler, M., Bolker, B., & Walker, S. (2014). Fitting Linear Mixed-Effects Models using lme4. *arXiv:1406.5823 [Stat]*. http://arxiv.org/abs/1406.5823

9. Bornkessel-Schlesewsky, I., & Schlesewsky, M. (2019). Toward a neurobiologically plausible model of language-related, negative event-related potentials. Frontiers in Psychology, 10(298), 1–17. 10.3389/fpsyg.2019.00298

10. Buzsáki, G., & Draguhn, A. (2004). Neuronal Oscillations in Cortical Networks. Science, 304(5679), 1926–1929. 10.1126/science.1099745

11. Buzsáki, G. (2005). Theta rhythm of navigation: Link between path integration and landmark navigation, episodic and semantic memory. Hippocampus, 15(7), 827–840. 10.1002/hipo.20113

12. Cavanagh, J. F., Figueroa, C. M., Cohen, M. X., & Frank, M. J. (2012). Frontal Theta Reflects Uncertainty and Unexpectedness during Exploration and Exploitation. Cerebral Cortex, 22(11), 2575–2586. 10.1093/cercor/bhr332

13. Cavanagh, J. F., & Frank, M. J. (2014). Frontal theta as a mechanism for cognitive control. Trends in Cognitive Sciences, 18(8), 414–421. 10.1016/j.tics.2014.04.012

14. Cavanagh, J. F., Frank, M. J., Klein, T. J., & Allen, J. J. B. (2010). Frontal theta links prediction errors to behavioral adaptation in reinforcement learning. NeuroImage, 49(4), 3198–3209. 10.1016/j.neuroimage.2009.11.080

15. Chen, Y. Y., & Caplan, J. B. (2017). Rhythmic Activity and Individual Variability in Recognition Memory: Theta Oscillations Correlate with Performance whereas Alpha Oscillations Correlate with ERPs. Journal of Cognitive Neuroscience, 29(1), 183–202. 10.1162/jocn_a_01033

16. Clark, A. (2013). Whatever next? Predictive brains, situated agents, and the future of cognitive science. Behavioral and Brain Sciences, 36(3), 181–204. 10.1017/S0140525X12000477

17. Cohen, M. X. (2011). Error-related medial frontal theta activity predicts cingulate-related structural connectivity. NeuroImage, 55(3), 1373–1383. 10.1016/j.neuroimage.2010.12.072

18. Collingridge, G. L., Peineau, S., Howland, J. G., & Wang, Y. T. (2010). Long-term depression in the CNS. Nature Reviews Neuroscience, 11(7), 459–473. 10.1038/nrn2867

19. Corcoran, A. W., Alday, P. M., Schlesewsky, M., & Bornkessel-Schlesewsky, I. (2018). Toward a reliable, automated method of individual alpha frequency (IAF) quantification. Psychophysiology, 55(7), e13064. 10.1111/psyp.13064

20. DeLong, K. A., Urbach, T. P., & Kutas, M. (2005). Probabilistic word pre-activation during language comprehension inferred from electrical brain activity. Nature Neuroscience, 8(8), Article 8. 10.1038/nn1504

21. den Ouden, H. E., Kok, P., & De Lange, F. P. (2012). How Prediction Errors Shape Perception, Attention, and Motivation. Frontiers in Psychology, 3. 10.3389/fpsyg.2012.00548

22. Dini, H., Simonetti, A., Bigne, E., & Bruni, L. E. (2022). EEG theta and N400 responses to congruent versus incongruent brand logos. Scientific Reports, 12(1), Article 1. 10.1038/s41598-022-08363-1

23. Donoghue, T., Dominguez, J., & Voytek, B. (2020). Electrophysiological Frequency Band Ratio Measures Conflate Periodic and Aperiodic Neural Activity. bioRxiv, 2020.01.11.900977. 10.1101/2020.01.11.900977

24. Donoghue, T., Haller, M., Peterson, E. J., Varma, P., Sebastian, P., Gao, R., Noto, T., Lara, A. H., Wallis, J. D., Knight, R. T., Shestyuk, A., & Voytek, B. (2020). Parameterizing neural power spectra into periodic and aperiodic components. Nature Neuroscience, 23(12), 1655–1665. 10.1038/s41593-020-00744-x

25. Eddine, S. N., Brothers, T., Wang, L., Spratling, M., & Kuperberg, G. R. (2024). A predictive coding model of the N400. Cognition, 246, 105755. 10.1016/j.cognition.2024.105755

26. Ergo, K., De Loof, E., Janssens, C., & Verguts, T. (2019). Oscillatory signatures of reward prediction errors in declarative learning. NeuroImage, 186, 137–145. 10.1016/j.neuroimage.2018.10.083

27. Exton-McGuinness, M. T. J., Lee, J. L. C., & Reichelt, A. C. (2015). Updating memories— The role of prediction errors in memory reconsolidation. Behavioural Brain Research, 278, 375–384. 10.1016/j.bbr.2014.10.011

28. Federmeier, K. D., & Kutas, M. (1999). A Rose by Any Other Name: Long-Term Memory Structure and Sentence Processing. Journal of Memory and Language, 41(4), 469–495. 10.1006/jmla.1999.2660

29. Federmeier, K. D., Wlotko, E. W., De Ochoa-Dewald, E., & Kutas, M. (2007). Multiple effects of sentential constraint on word processing. Brain Research, 1146, 75–84. 10.1016/j.brainres.2006.06.101

30. Feldman, H., & Friston, K. J. (2010). Attention, Uncertainty, and Free-Energy. Frontiers in Human Neuroscience, 4. 10.3389/fnhum.2010.00215

31. Fernández, G., Weyerts, H., Tendolkar, I., Smid, H. G. O. M., Scholz, M., & Heinze, H. J. (1998). Event-related potentials of verbal encoding into episodic memory: Dissociation between the effects of subsequent memory performance and distinctiveness. Psychophysiology, 35(6), 709–720. 10.1111/1469-8986.3560709

32. Fernández, R. S., Boccia, M. M., & Pedreira, M. E. (2016). The fate of memory: Reconsolidation and the case of Prediction Error. Neuroscience & Biobehavioral Reviews, 68, 423–441. 10.1016/j.neubiorev.2016.06.004

33. Fitz, H., & Chang, F. (2019). Language ERPs reflect learning through prediction error propagation. Cognitive Psychology, 111, 15–52. 10.1016/j.cogpsych.2019.03.002

34. Fox, J., & Weisberg, S. (2019). An R Companion to Applied Regression. SAGE Publications.

34a. Friston, K. (2005). A theory of cortical responses. Philosophical Transactions of the Royal Society B: Biological Sciences, 360(1456), 815–836. 10.1098/rstb.2005.1622

35. Gramfort, A., Luessi, M., Larson, E., Engemann, D. A., Strohmeier, D., Brodbeck, C., Goj, R., Jas, M., Brooks, T., Parkkonen, L., & Hämäläinen, M. (2013). MEG and EEG data analysis with MNE-Python. Frontiers in Neuroscience, 7(7), 267–267. 10.3389/fnins.2013.00267

36. Grandy, T. H., Werkle-Bergner, M., Chicherio, C., Lövdén, M., Schmiedek, F., & Lindenberger, U. (2013). Individual alpha peak frequency is related to latent factors of general cognitive abilities. NeuroImage, 79, 10–18. 10.1016/j.neuroimage.2013.04.059

37. Greve, A., Cooper, E., Kaula, A., Anderson, M. C., & Henson, R. (2017). Does prediction error drive one-shot declarative learning? Journal of Memory and Language, 94, 149–165. 10.1016/j.jml.2016.11.001

38. Gronau, N., & Shachar, M. (2015). Contextual consistency facilitates long-term memory of perceptual detail in barely seen images. Journal of Experimental Psychology: Human Perception and Performance, 41(4), 1095–1111. 10.1037/xhp0000071

39. Hald, L. A., Bastiaansen, M. C. M., & Hagoort, P. (2006). EEG theta and gamma responses to semantic violations in online sentence processing. Brain and Language, 96(1), 90–105. 10.1016/j.bandl.2005.06.007

40. Hanslmayr, S., Spitzer, B., & Bauml, K. H. (2009). Brain Oscillations Dissociate between Semantic and Nonsemantic Encoding of Episodic Memories. Cerebral Cortex, 19(7), 1631–1640. 10.1093/cercor/bhn197

41. Hanslmayr, S., & Staudigl, T. (2014). How brain oscillations form memories—A processing based perspective on oscillatory subsequent memory effects. NeuroImage, 85, 648– 655. 10.1016/j.neuroimage.2013.05.121

42. Hanslmayr, S., Staudigl, T., & Fellner, M. C. (2012). Oscillatory power decreases and long-term memory: The information via desynchronization hypothesis. Frontiers in Human Neuroscience, 6. 10.3389/fnhum.2012.00074

43. Hasselmo, M. E. (2005). What is the function of hippocampal theta rhythm?—Linking behavioral data to phasic properties of field potential and unit recording data. Hippocampus, 15(7), 936–949. 10.1002/hipo.20116

44. Hasselmo, M. E., & Stern, C. E. (2014). Theta rhythm and the encoding and retrieval of space and time. NeuroImage, 85, 656–666. 10.1016/j.neuroimage.2013.06.022

45. He, B. J. (2014). Scale-free brain activity: Past, present, and future. Trends in Cognitive Sciences, 18(9), 480–487. 10.1016/j.tics.2014.04.003

46. He, B. J., Zempel, J. M., Snyder, A. Z., & Raichle, M. E. (2010). The Temporal Structures and Functional Significance of Scale-free Brain Activity. Neuron, 66(3), 353–369. 10.1016/j.neuron.2010.04.020

47. Hodapp, A., & Rabovsky, M. (2021). The N400 ERP component reflects an error-based implicit learning signal during language comprehension. European Journal of Neuroscience, 54(9), 7125–7140. 10.1111/ejn.15462

48. Hohwy, J. (2020). New directions in predictive processing. Mind & Language, 35(2), 209–223. 10.1111/mila.12281

49. Holroyd, C. B., & Coles, M. G. H. (2002). The neural basis of human error processing: Reinforcement learning, dopamine, and the error-related negativity. Psychological Review, 109(4), 679–709. 10.1037/0033-295X.109.4.679

50. Hsieh, L.-T., & Ranganath, C. (2014). Frontal midline theta oscillations during working memory maintenance and episodic encoding and retrieval. NeuroImage, 85, 721–729. 10.1016/j.neuroimage.2013.08.003

51. Huang, Y., & Rao, R. P. N. (2011). Predictive coding. WIREs Cognitive Science, 2(5), 580–593. 10.1002/wcs.142

52. Hubbard, R. J., & Federmeier, K. D. (2024). The Impact of Linguistic Prediction Violations on Downstream Recognition Memory and Sentence Recall. Journal of Cognitive Neuroscience, 36(1), 1–23. 10.1162/jocn_a_02078

53. Hubbard, R. J., Rommers, J., Jacobs, C. L., & Federmeier, K. D. (2019). Downstream Behavioral and Electrophysiological Consequences of Word Prediction on Recognition Memory. Frontiers in Human Neuroscience, 13. https://www.frontiersin.org/articles/10.3389/fnhum.2019.00291

54. Jano, S., Chatburn, A., Cross, Z., Schlesewsky, M., & Bornkessel-Schlesewsky, I. (2024). How predictability and individual alpha frequency shape memory: Insights from an event-related potential investigation. Neurobiology of Learning and Memory, 108006. 10.1016/j.nlm.2024.108006

55. Jano, S., Cross, Z. R., Chatburn, A., Schlesewsky, M., & Bornkessel-Schlesewsky, I. (2024). Prior Context and Individual Alpha Frequency Influence Predictive Processing during Language Comprehension. Journal of Cognitive Neuroscience, 1–39. 10.1162/jocn_a_02196

56. Jano, S., Romeo, J., Hendrickx, M. D., Schlesewsky, M., & Chatburn, A. (2021). Sleep influences neural representations of true and false memories: An event-related potential study. Neurobiology of Learning and Memory, 186, 107553. 10.1016/j.nlm.2021.107553

57. Jas, M., Engemann, D., Raimondo, F., Bekhti, yousra, & Gramfort, A. (2016, June). Automated rejection and repair of bad trials in MEG/EEG. 6th International Workshop on Pattern Recognition in Neuroimaging (PRNI). https://hal.archives-ouvertes.fr/hal-01313458

58. Klimesch, W. (1997). EEG-alpha rhythms and memory processes. International Journal of Psychophysiology, 26(1), 319–340. 10.1016/S0167-8760(97)00773-3

59. Klimesch, W. (1999). EEG alpha and theta oscillations reflect cognitive and memory performance: A review and analysis. Brain Research. Brain Research Reviews, 29(2– 3), 169–195. 10.1016/s0165-0173(98)00056-3

60. Klimesch, W., Doppelmayr, M., Russegger, H., & Pachinger, T. (1996). Theta band power in the human scalp EEG and the encoding of new information. NeuroReport, 7(7), 1235. 10.1097/00001756-199605170-00002

61. Klimesch, W., Doppelmayr, M., Schimke, H., & Ripper, B. (1997). Theta synchronization and alpha desynchronization in a memory task. Psychophysiology, 34(2), 169–176. 10.1111/j.1469-8986.1997.tb02128.x

62. Klimesch, W., Sauseng, P., & Hanslmayr, S. (2007). EEG alpha oscillations: The inhibition– timing hypothesis. Brain Research Reviews, 53(1), 63–88. 10.1016/j.brainresrev.2006.06.003

63. Klimesch, W., Schimke, H., Doppelmayr, M., Ripper, B., Schwaiger, J., & Pfurtscheller, G. (1996). Event-related desynchronization (ERD) and the Dm effect: Does alpha desynchronization during encoding predict later recall performance? International Journal of Psychophysiology, 24(1), 47–60. 10.1016/S0167-8760(96)00054-2

64. Klimesch, W., Schimke, H., Ladurner, G., & Pfurtscheller, G. (1990). Alpha frequency and memory performance. Journal of Psychophysiology, 4(4), 381–390.

65. Klimesch, W., Schimke, H., & Pfurtscheller, G. (1993). Alpha frequency, cognitive load and memory performance. Brain Topography, 5(3), 241–251. 10.1007/BF01128991

66. Kutas, M., & Federmeier, K. D. (2011). Thirty Years and Counting: Finding Meaning in the N400 Component of the Event-Related Brain Potential (ERP). Annual Review of Psychology, 62(1), 621–647. 10.1146/annurev.psych.093008.131123

67. Kutas, M., & Hillyard, S. A. (1980). Reading Senseless Sentences: Brain Potentials Reflect Semantic Incongruity. Science, 207(4427), 203–205. 10.1126/science.7350657

68. Kutas, M., & Hillyard, S. A. (1984). Brain potentials during reading reflect word expectancy and semantic association. Nature, 307(5947), Article 5947. 10.1038/307161a0

69. Leung, L. S., & Law, C. S. H. (2020). Phasic modulation of hippocampal synaptic plasticity by theta rhythm. Behavioral Neuroscience, 134(6), 595–612. 10.1037/bne0000354

70. Long, N. M., Burke, J. F., & Kahana, M. J. (2014). Subsequent memory effect in intracranial and scalp EEG. NeuroImage, 84, 488–494. 10.1016/j.neuroimage.2013.08.052

71. Luck, S. J. (2014). An introduction to the event-related potential technique (Second edition). The MIT Press.

72. Lüdecke, D. (2018). ggeffects: Tidy Data Frames of Marginal Effects from Regression Models. Journal of Open Source Software, 3(26), 772. 10.21105/joss.00772

73. Luu, P., Tucker, D. M., & Makeig, S. (2004). Frontal midline theta and the error-related negativity: Neurophysiological mechanisms of action regulation. Clinical Neurophysiology, 115(8), 1821–1835. 10.1016/j.clinph.2004.03.031

74. Mack, M. L., Love, B. C., & Preston, A. R. (2018). Building concepts one episode at a time: The hippocampus and concept formation. Neuroscience Letters, 680, 31–38. 10.1016/j.neulet.2017.07.061

75. Martin, S. J., Grimwood, P. D., & Morris, R. G. M. (2000). Synaptic Plasticity and Memory: An Evaluation of the Hypothesis. Annual Review of Neuroscience, 23(Volume 23, 2000), 649–711. 10.1146/annurev.neuro.23.1.649

76. Mas-Herrero, E., & Marco-Pallarés, J. (2014). Frontal Theta Oscillatory Activity Is a Common Mechanism for the Computation of Unexpected Outcomes and Learning Rate. Journal of Cognitive Neuroscience, 26(3), 447–458. 10.1162/jocn_a_00516

77. McNicol, D. (2004). A Primer of Signal Detection Theory. Taylor & Francis Group. http://ebookcentral.proquest.com/lib/unisa/detail.action?docID=227463

78. Meeuwissen, E. B., Takashima, A., Fernández, G., & Jensen, O. (2011). Increase in posterior alpha activity during rehearsal predicts successful long-term memory formation of word sequences. Human Brain Mapping, 32(12), 2045–2053. 10.1002/hbm.21167

79. Mumford, D. (1992). On the computational architecture of the neocortex. Biological Cybernetics, 66(3), 241–251. 10.1007/BF00198477

80. Munneke, G.-J., Nap, T. S., Schippers, E. E., & Cohen, M. X. (2015). A statistical comparison of EEG time- and time–frequency domain representations of error processing. Brain Research, 1618, 222–230. 10.1016/j.brainres.2015.05.030

81. Ordin, M., Polyanskaya, L., Soto, D., & Molinaro, N. (2020). Electrophysiology of statistical learning: Exploring the online learning process and offline learning product. European Journal of Neuroscience, 51(9), 2008–2022. 10.1111/ejn.14657

82. Ortiz-Tudela, J., Nolden, S., Pupillo, F., Ehrlich, I., Schommartz, I., Turan, G., & Shing, Y. L. (2023). Not what u expect: Effects of prediction errors on item memory. Journal of Experimental Psychology: General, 152(8), 2160–2176. 10.1037/xge0001367

83. Pratt, H. (2011). Sensory ERP components. The Oxford handbook of event-related potential components, 89–114.

84. Pu, Y., Cheyne, D., Sun, Y., & Johnson, B. W. (2020). Theta oscillations support the interface between language and memory. NeuroImage, 215, 116782. 10.1016/j.neuroimage.2020.116782

85. Rabovsky, M., & McRae, K. (2014). Simulating the N400 ERP component as semantic network error: Insights from a feature-based connectionist attractor model of word meaning. Cognition, 132(1), 68–89. 10.1016/j.cognition.2014.03.010

86. Rao, R. P. N., & Ballard, D. H. (1999). Predictive coding in the visual cortex: A functional interpretation of some extra-classical receptive-field effects. Nature Neuroscience, 2(1). 10.1038/4580

87. Rauss, K., & Born, J. (2017). A Role of Sleep in Forming Predictive Codes. In N. Axmacher & B. Rasch (Eds.), Cognitive Neuroscience of Memory Consolidation (pp. 117–132). Springer International Publishing. 10.1007/978-3-319-45066-7_8

88. Rouhani, N., Norman, K. A., & Niv, Y. (2018). Dissociable effects of surprising rewards on learning and memory. *Journal of Experimental Psychology: Learning*, Memory, and Cognition, 44(9), 1430–1443. 10.1037/xlm0000518

89. Samaha, J., & Postle, B. R. (2015). The Speed of Alpha-Band Oscillations Predicts the Temporal Resolution of Visual Perception. Current Biology, 25(22), 2985–2990. 10.1016/j.cub.2015.10.007

90. Schacter, D. L., Addis, D. R., & Buckner, R. L. (2008). Episodic Simulation of Future Events. Annals of the New York Academy of Sciences, 1124(1), 39–60. 10.1196/annals.1440.001

91. Schnitzler, A., & Gross, J. (2005). Normal and pathological oscillatory communication in the brain. Nature Reviews Neuroscience, 6(4), 285–296. 10.1038/nrn1650

92. Sederberg, P. B., Kahana, M. J., Howard, M. W., Donner, E. J., & Madsen, J. R. (2003). Theta and Gamma Oscillations during Encoding Predict Subsequent Recall. The Journal of Neuroscience, 23(34), 10809–10814. 10.1523/JNEUROSCI.23-34-10809.2003

93. Sherman, B. E., Graves, K. N., Huberdeau, D. M., Quraishi, I. H., Damisah, E. C., & Turk-Browne, N. B. (2022). Temporal Dynamics of Competition between Statistical Learning and Episodic Memory in Intracranial Recordings of Human Visual Cortex. Journal of Neuroscience, 42(48), 9053–9068. 10.1523/JNEUROSCI.0708-22.2022

94. Sherman, B. E., & Turk-Browne, N. B. (2020). Statistical prediction of the future impairs episodic encoding of the present. Proceedings of the National Academy of Sciences, 117(37), 22760–22770. 10.1073/pnas.2013291117

95. Sinclair, A. H., Manalili, G. M., Brunec, I. K., Adcock, R. A., & Barense, M. D. (2021). Prediction errors disrupt hippocampal representations and update episodic memories. Proceedings of the National Academy of Sciences, 118(51), e2117625118. 10.1073/pnas.2117625118

96. Sitnikova, T., Holcomb, P. J., Kiyonaga, K. A., & Kuperberg, G. R. (2008). Two Neurocognitive Mechanisms of Semantic Integration during the Comprehension of Visual Real-world Events. Journal of Cognitive Neuroscience, 20(11), 2037–2057. 10.1162/jocn.2008.20143

97. Trujillo, L. T., & Allen, J. J. B. (2007). Theta EEG dynamics of the error-related negativity. Clinical Neurophysiology, 118(3), 645–668. 10.1016/j.clinph.2006.11.009

98. Turan, G., Ehrlich, I., Shing, Y. L., & Nolden, S. (2023). From generating to violating predictions: The effects of prediction error on episodic memory. PsyArXiv. 10.31234/osf.io/zm29a

99. Urgen, B. A., Kutas, M., & Saygin, A. P. (2018). Uncanny valley as a window into predictive processing in the social brain. Neuropsychologia, 114, 181–185. 10.1016/j.neuropsychologia.2018.04.027

100. Vallat, R., & Walker, M. P. (2021). An open-source, high-performance tool for automated sleep staging. Elife, 10, e70092. 10.7554/eLife.70092

101. Van Berkum, J. J. A. (2010). The brain is a prediction machine that cares about good and bad—Any implications for neuropragmatics? Italian Journal of Linguistics, 22, 181–208. 10/component/file_539546/vanberkum-iljpap2010-definitive.pdf

102. Van Berkum, J. J. A., Zwitserlood, P., Hagoort, P., & Brown, C. M. (2003). When and how do listeners relate a sentence to the wider discourse? Evidence from the N400 effect. Cognitive Brain Research, 17(3), 701–718. 10.1016/S0926-6410(03)00196-4

103. Van de Vijver, I., Ridderinkhof, K. R., & Cohen, M. X. (2011). Frontal Oscillatory Dynamics Predict Feedback Learning and Action Adjustment. Journal of Cognitive Neuroscience, 23(12), 4106–4121. 10.1162/jocn_a_00110

104. Wen, H., & Liu, Z. (2016). Separating Fractal and Oscillatory Components in the Power Spectrum of Neurophysiological Signal. Brain Topography, 29(1), 13–26. 10.1007/s10548-015-0448-0

105. White, T. P., Jansen, M., Doege, K., Mullinger, K. J., Park, S. B., Liddle, E. B., Gowland, P. A., Francis, S. T., Bowtell, R., & Liddle, Peter. F. (2013). Theta power during encoding predicts subsequent-memory performance and default mode network deactivation. Human Brain Mapping, 34(11), 2929–2943. 10.1002/hbm.22114

106. Wickham, H. (2016). Ggplot2 citation info. Retrieved March 25, 2021, from https://cran.r-project.org/web/packages/ggplot2/citation.html

107. Worthen, J. B., & Roark, B. (2002). Free Recall Accuracy for Common and Bizarre Verbal Information. The American Journal of Psychology, 115(3), 377–394. 10.2307/1423423

